# Epistatic Networks Associated with Parent-of-Origin Effects on Metabolic Traits

**DOI:** 10.1101/579748

**Authors:** Juan F Macias-Velasco, Celine L. St. Pierre, Jessica P Wayhart, Li Yin, Larry Spears, Mario A. Miranda, Katsuhiko Funai, James M Cheverud, Clay F Semenkovich, Heather A Lawson

## Abstract

Parent-of-origin effects (POE) are unexpectedly common in complex traits, including metabolic and neurological diseases. POE can also be modified by the environment, but the architecture of these gene-by-environmental effects on phenotypes remains to be unraveled. Previously, quantitative trait loci (QTL) showing context-specific POE on metabolic traits were mapped in the F_16_ generation of an advanced intercross between LG/J and SM/J inbred mice. However, these QTL were not enriched for known imprinted genes, suggesting another mechanism is needed to explain these POE phenomena. Here, we use a simple yet powerful F_1_ reciprocal cross model to test the hypothesis that non-imprinted genes can generate complex POE on metabolic traits through genetic interactions with imprinted genes. Male and female mice from a F_1_ reciprocal cross of LG/J and SM/J strains were fed either high or low fat diets. We generated expression profiles from three metabolically-relevant tissues: hypothalamus, white adipose, and liver. We identified two classes of parent-of-origin expression biases: genes showing parent-of-origin-dependent allele-specific expression and biallelic genes that are differentially expressed by reciprocal cross. POE patterns of both gene classes are highly tissue-and context-specific, sometimes occurring only in one sex and/or diet cohort in a particular tissue. We then constructed tissue-specific interaction networks among genes from these two classes of POE. A key subset of gene pairs show significant epistasis in the F_16_ LG/J x SM/J advanced intercross data in cases where the biallelic gene fell within a previously-identified metabolic POE QTL interval. We highlight one such interaction in adipose, between *Nnat* and *Mogat1*, which associates with POE on multiple adiposity traits. Both genes localize to the endoplasmic reticulum of adipocytes and play a role in adipogenesis. Additionally, expression of both genes is significantly correlated in human visceral adipose tissue. The genes and networks we present here represent a set of actionable interacting candidates that can be probed to further identify the machinery driving POE on complex traits.

---

Parent-of-origin effects, where an allele’s phenotypic effect depends on whether it is inherited maternally or paternally, are associated with a wide range of common complex traits and diseases^1^. Several mechanisms can cause parent-of-origin effects on phenotype including genomic imprinting, maternal/paternal effects, and sex-biased gene-specific trinucleotide expansions^2–4^. The best characterized parent-of-origin effect on phenotype is genomic imprinting, an epigenetic process in which either the maternally or paternally inherited allele is silenced. Parent-of-origin effects are often associated with diseases related to metabolism, neurological function, or both. Metabolic diseases include transient neonatal diabetes^5–8^, type-1 diabetes^9^, type-2 diabetes^10,11^, some cancers^12,13^, metabolic syndrome, Beckwith-Wiedemann syndrome, Wilm’s tumors^14^, insulinomas^12^, Silver–Russell syndrome^15,16^, and variants of Albright’s hereditary osteodystrophy^17,18^. Neurological diseases include Alzheimers^19–21^, myoclonus-dystonia syndrome^22^, and Jervell and Lange-Nielsen syndrome^23,24^. Diseases that are both metabolic and neurological in nature include Prader-Willi and Angelman syndromes^25,26^. Parent-of-origin effects associated with disease can exhibit variance among individuals, and may be modified by the environment^1,27–29^. Unraveling the genetic architecture of these effects will improve efforts to predict phenotype from (epi)genotype. This can direct research aimed at developing novel therapeutic strategies for diseases associated with parent-of-origin imprinted effects^30^.

Parent-of-origin effects on complex traits and disease are unexpectedly common, despite there being few known imprinted genes, suggesting that canonical imprinting mechanisms are not sufficient to account for these phenomena^31,32^. We hypothesize that interactions between imprinted genes and non-imprinted genes with equivalent expression of parental alleles (biallelic genes), may underlie some of these effects on phenotype (**Figure 1**). These interactions may be the means by which parent-of-origin signals can propagate from epigenetic marks to gross phenotypic variation, despite the relative rarity of canonically imprinted genes.

**Figure 1.**
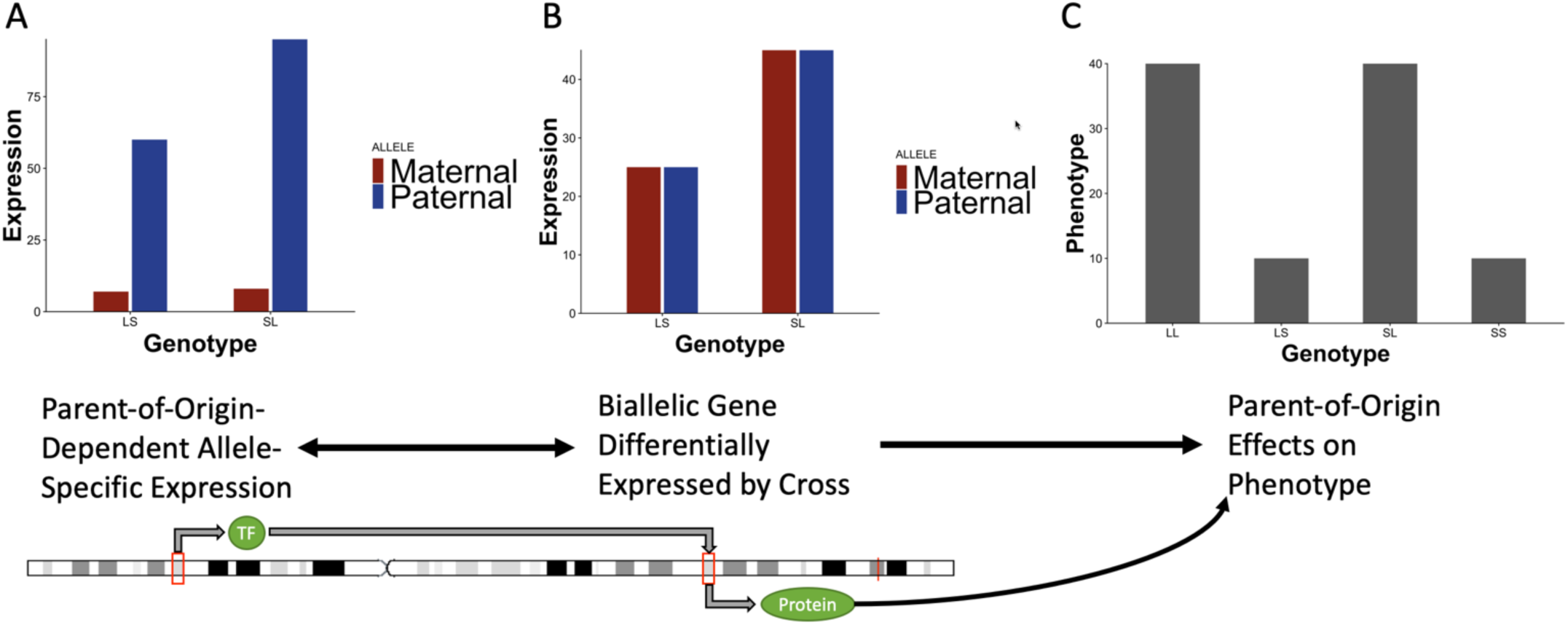
Proposed mechanism for how genes showing parent-of-origin-dependent allele -specific expression interacting with biallelic genes can lead to parent-of-origin effects on phenotype. **A.** A gene with parent-of-origin-dependent allele-specific expression shows differential expression based on allelic identity (L or S). This gene may be a transcription factor that binds to the promotor of a biallelic gene. **B.** The differential expression of the gene with parent-of-origin-dependent allele-specific expression, or additional variants in the promotor region affecting binding efficiency, leads to asymmetry in expression between reciprocal heterozygotes. **C.** This results in a parent-of-origin effect on phenotype. Alleles are ordered maternal | paternal.

Interactions among imprinted genes are likely, as they tend to be clustered together on the genome and are significantly co-expressed^33,34^. Network analysis of imprinted genes indicates that they are particularly “interactive”, and are enriched in complex networks that include both imprinted and non-imprinted genes^29,35,36^. Genetic interactions between imprinted genes and non-imprinted genes have already been shown to contribute to variation in metabolic phenotypes. For example, the maternally expressed transcription factor KLF14 (kruppel-like factor 14) has been shown to regulate biallelic genes related to adiposity^10,16^. Mapping studies have identified two SNPs upstream of KLF14 that are associated with type-2 diabetes and cholesterol levels^37,38^. Both variants have maternally-restricted *cis*-regulatory associations with KLF14 expression in adipose tissue^39^. eQTL analyses in human data found that one of the SNPs is enriched for *trans*-associations with KLF family transcription factor binding sites in subcutaneous white adipose tissue, indicating that KLF14 may be a master transcriptional regulator in adipose^10^. Together, this suggests that the imprinted gene KLF14 propagates a parent-of-origin effect on non-imprinted (biallelic) genes that contribute to variation in gene expression and, ultimately, to variation in adiposity traits. Whether additional pairs of imprinted and biallelic genes are similarly co-expressed and whether these parent-of-origin dependent network patterns transfer to other complex traits (metabolic or not) remain important open questions. The transience of these gene networks across tissues or environmental contexts is also understudied. Interactions between imprinted and biallelic genes could explain some of the observed parent-of-origin effect patterns associated with regions lacking obvious candidate genes, as described in a recent survey of 97 complex traits measured in outbred mice^29^.

In this study, we use a simple yet powerful F_1_ reciprocal cross model of the LG/J and SM/J inbred mice (LxS and SxL) to test the hypothesis that non-imprinted genes can generate complex parent-of-origin effects on dietary-obesity phenotypes through genetic interactions with imprinted genes. LG/J and SM/J are frequently used in metabolic studies because these strains vary in their metabolic response to dietary fat^40^. Quantitative trait loci (QTL) showing parent-of-origin effects on dietary obesity traits have been mapped in the F_16_ generation of an advanced intercross between LG/J and SM/J^41–44^. Most of these QTL had additive effects, but ≈60% of them also showed parent-of-origin effects in some sex and/or dietary context. However, permutation analyses reveal that these parent-of-origin effect QTL are not enriched for known imprinted genes (**Supplemental Figure 1**).

We generated expression profiles from 20 week-old F_1_ animals in order to match the age of the F_16_ LG/J x SM/J advanced intercross population in which dietary-obesity QTL having parent-of-origin effects were mapped. F_1_ reciprocal cross (LxS and SxL) animals were subject to the same high and low fat diets and phenotyping protocols as the previously-studied F_16_ animals to keep environmental contexts consistent. We identify genes showing parent-of-origin-dependent allele-specific expression in three metabolically-relevant tissues: hypothalamus, white adipose (reproductive fatpad), and liver. Some of these genes are canonically imprinted, but most have an allele that is partially silenced, generating a parent-of-origin-dependent expression bias. We characterize interactions among these genes and biallelic genes that are differentially expressed by reciprocal cross under different sex and/or diet contexts. Further, we test for epistasis between interacting gene pairs where the biallelic gene is located within a previously-mapped dietary-obesity QTL showing parent-of-origin effects in the F_16_ population.

## RESULTS

### Parent-of-origin-dependent allele-specific expression shows a high degree of tissue-specificity

We identified genes with significant parent-of-origin-dependent allele-specific expression in hypothalamus (n=108), white adipose (n=102), and liver (n=109) tissues across sex and dietary contexts (**Supplemental Table 1**). We calculated parent-of-origin effect (POE) scores for each expressed gene by subtracting the mean L_bias_ of the SxL (maternal SM/J, paternal LG/J) cross from the mean L_bias_ of the LxS (maternal LG/J, paternal SM/J) cross of all mice in a certain diet-by-sex context (high or low fat diets, equal representation of males and females). POE scores range from completely maternally-expressed (−1), to biallelic (0), to completely paternally expressed (+1). A total of 298 unique genes show parent-of-origin-dependent allele-specific expression across these tissues. We also observe a relationship between sex and dietary context-and tissue-dependency among their expression profiles. If a gene shows parent-of-origin-dependent allele-specific expression across all diet-by-sex contexts in a given tissue (context-independent), then it tends to show parent-of-origin allele-specific expression across all contexts in the other two tissues (tissue-independent). However, if a gene shows varying degrees of parent-of-origin-dependent allele-specific expression across diet-by-sex contexts in a given tissue (context-dependency), then it also tends to have variable parent-of-origin-dependent allele-specific expression patterns across these contexts in the other two tissues (tissue-dependency) (**Figure 2**). Canonically imprinted genes are the most likely to be both context-and tissue-independent.

**Figure 2.**
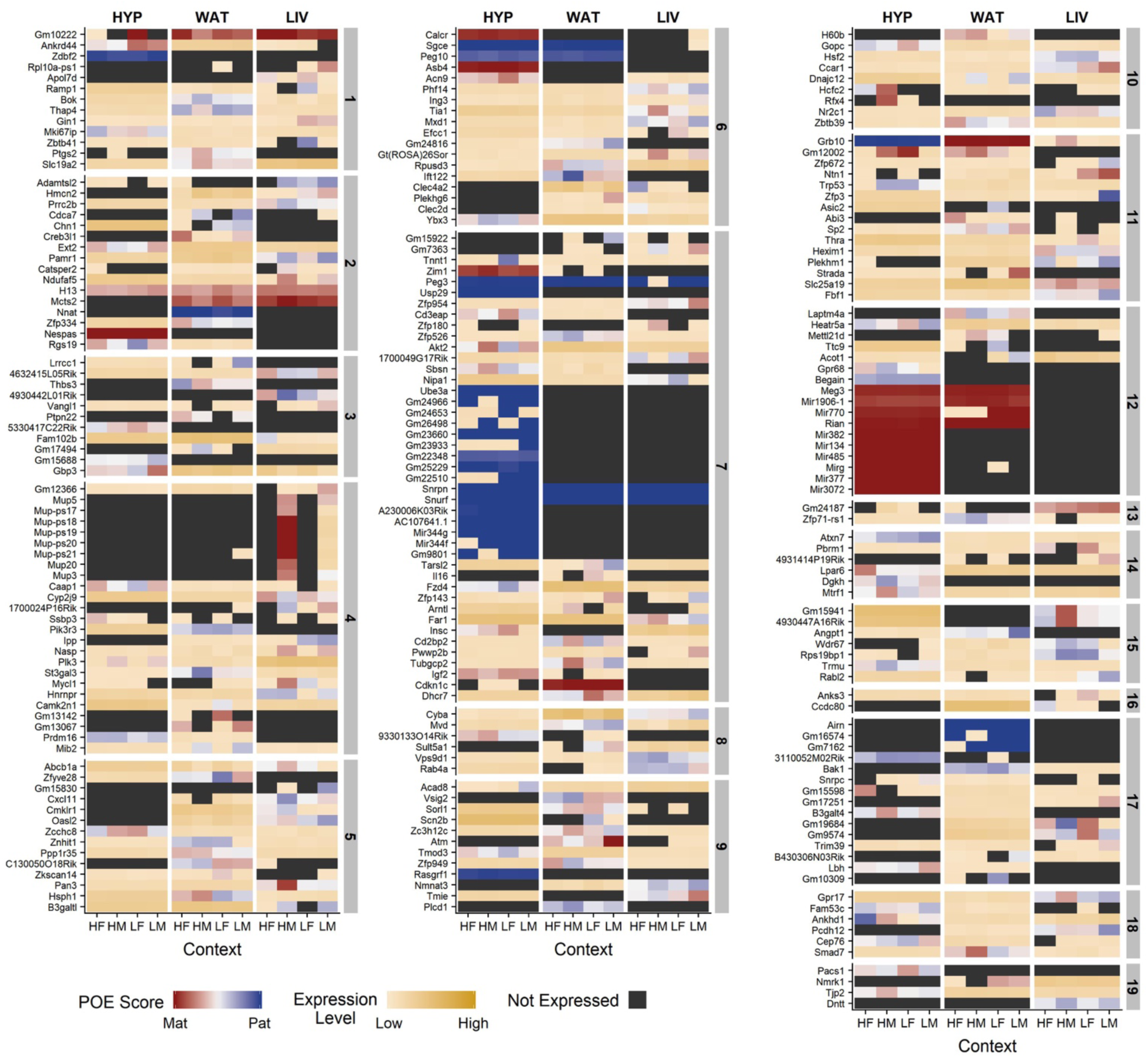
Genes show parent-of-origin dependent allele-specific expression in both context-dependent and context-independent manners. An expression profile heat map of genes showing significant parent-of-origin dependent allele-specific expression in at least one diet-by**-**sex context across three metabolically-relevant tissues. Expression levels range from white (lowly expressed) to dark goldenrod (highly expressed). If a gene is not expressed in a certain tissue and diet-by-sex context, then it is shown as black. Parent-of-origin (POE) scores range from completely maternally-expressed (−1, Mat, dark red), to biallelic (0, white), to completely paternally-expressed (+1, Pat, dark blue). The x-axis denotes diet-by-sex context (HF = high fat-fed females; HM = high fat-fed males; LF = low fat-fed females; LM = low fat-fed males) across three tissues (HYP = hypothalamus; WAT = white adipose; LIV = liver). The y-axis denotes genes with significant parent-of-origin dependent allele-specific expression ordered by genomic position across the chromosomes (vertical grey panels).

### Parent-of-origin-dependent expression biases are context-specific

Parent-of-origin-dependent expression biases are highly context-specific in both genes showing allele-specific expression and biallelic genes that are differentially expressed by cross (**Figure 3**; **Supplemental Tables 1 and 2**). For allele-specific expression, context-specificity was assigned by the most complex model term for which a significant parent-of-origin-dependent expression bias was observed. For biallelic genes differentially expressed by cross, the significance of the nested terms (**see Methods**) was compared to identify the model term that produced the most significant differential expression. More biallelic genes that are differentially expressed by cross are context-specific than genes showing parent-of-origin-dependent allele-specific expression. However, the degree of context-specificity varies by tissue; hypothalamus has the lowest proportion of context-specific parent-of-origin-dependent allele-specific expression (hypothalamus = 44.78%; white adipose = 48.7%; liver = 49.54%) as well as the highest proportion of biallelic genes that are differentially expressed by cross (hypothalamus = 95.16%; white adipose = 92.25%; liver = 94.1%).

**Figure 3.**
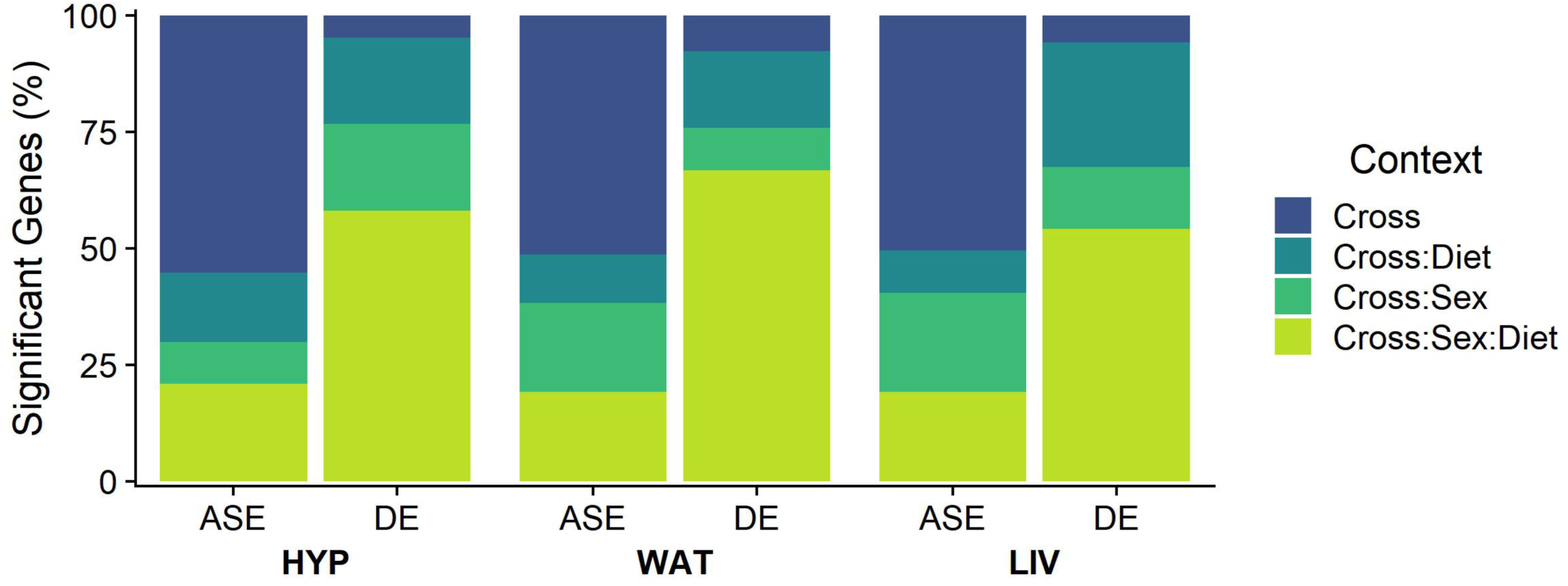
Parent-of-origin-dependent expression biases are context-specific. Parent-of-origin expression biases are considered significant if any of the Cross (Blue), Cross:Sex (Green), Cross:Diet (Teal), or Cross:Sex:Diet (lime green) terms were significant in the models tested. Parent-of-origin dependent biases show a high degree of context-specificity in both genes showing parent-of-origin-dependent expression biases and biallelic genes that are differentially expressed by cross. This context-specificity is also tissue-dependent. ASE = parent-of-origin-dependent allele-specific expression biases; DE = biallelic genes differentially expressed by cross (LxS or SxL); HYP = hypothalamus; WAT = white adipose; LIV = liver.

### Genes showing parent-of-origin effects form highly interconnected networks

Networks were constructed in each tissue from genes showing parent-of-origin-dependent allele-specific expression biases (hypothalamus = 109; white adipose = 102; liver = 108) and biallelic genes that are differentially expressed by cross (hypothalamus = 123; white adipose = 427; liver = 203). Hypothalamus had more genes that demonstrate both classes of parent-of-origin biases, allele-specific expression and differential expression by cross, than either white adipose or liver. However, many of those genes did not pass our significance thresholds and thus were not included in these network analyses (**Supplemental Table 3**). Interacting gene pairs were predicted by modeling the expression of biallelic genes that are significantly differentially expressed by cross as a function of the expression of genes showing significant parent-of-origin-dependent allele-specific expression, their allelic bias (L_bias_), diet, sex, and the diet-by-sex interaction. Genes showing parent-of-origin effects form highly interconnected networks in each tissue (**Figure 4**). Networks are comprised of 340 interacting gene pairs in hypothalamus, 2,570 in white adipose, and 1,171 in liver. These pairs represent both unique genes showing parent-of-origin-dependent allele-specific expression biases (hypothalamus = 83; white adipose = 82; liver = 86) and unique biallelic genes that are differentially expressed by cross (hypothalamus = 80; white adipose = 385; liver = 198) in some context (reciprocal cross, diet, sex, diet-by-sex).

**Figure 4.**
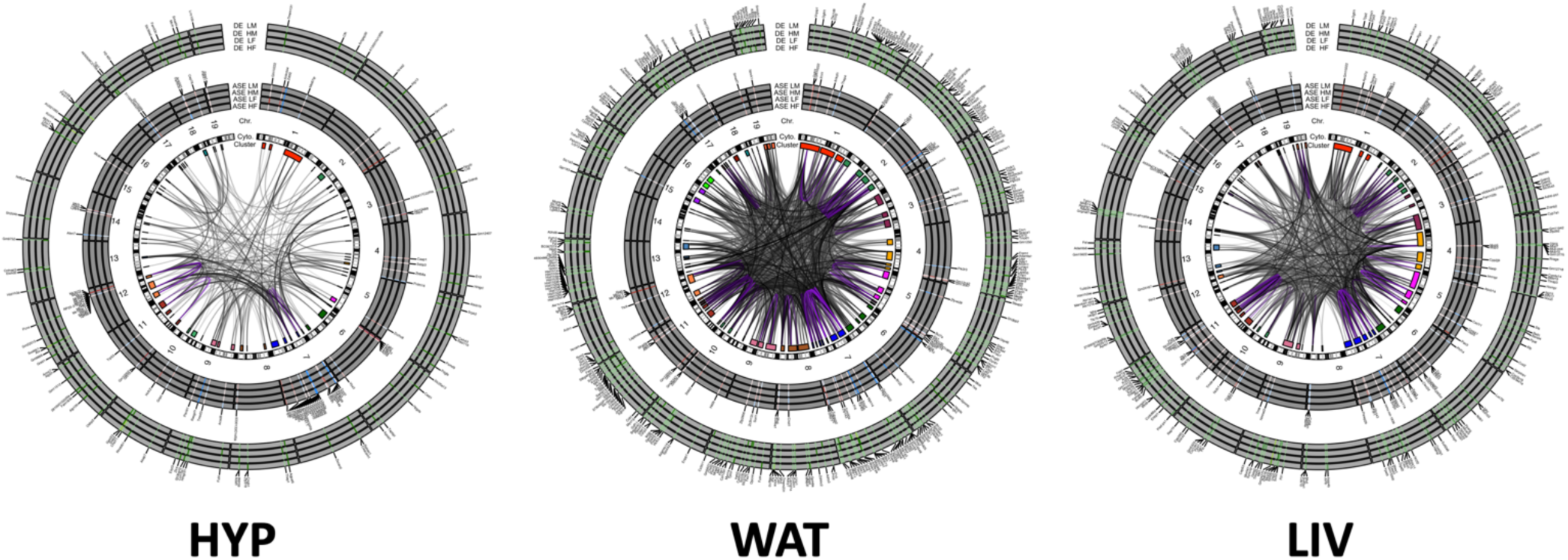
Genes showing parent-of-origin effects form highly interconnected networks. Networks are comprised of two classes of parent-of-origin effect: parent-of-origin-dependent allele-specific expression (middle tracks) and biallelic genes showing differential expression by cross, LxS or SxL (outer tracks). Genes showing parent-of-origin-dependent allele-specific expression are colored coded by their parent-of-origin effect score, with blue indicating a paternal bias and red indicating a maternal bias. Biallelic genes showing differential expression between crosses are colored by their absolute log_2_ fold change on a green gradient. Darker green indicates a larger fold change magnitude. The inner-most track represents the chromosome and the boxes represent clusters of genes. The clusters are color-coded by chromosome for ease of visualization. Significant interactions are denoted with a line connecting their genomic positions, with *cis*-interactions colored purple and *trans*-interactions colored black. HYP = hypothalamus; WAT = white adipose; LIV = liver.

Genes showing parent-of-origin-dependent allele-specific expression biases tend to cluster throughout the genome in a way that is comparable to how canonically imprinted genes are clustered. The degree of clustering varies by tissue, however, and hypothalamus shows the strongest clustering of genes. This is consistent with our finding that the hypothalamus has the highest proportion of genes showing context-independent parent-of-origin-dependent allele specific expression biases as well as the highest proportion of expressed canonically imprinted genes of the three tissues we assayed. Most significant interactions between genes showing parent-of-origin-dependent allele-specific expression biases and biallelic genes differentially expressed by cross are trans-chromosomal. This indicates that we are not identifying covariation among pairs that are controlled by the same regulatory element, as is often found among canonically imprinted genes that are controlled by the same *cis*-regulatory machinery^34,45^.

Enrichment analyses of significantly interacting genes reveals over-representation of canonically imprinted genes in hypothalamus, over-representation of genes involved in regulatory activity and iron metabolism in liver, and over-representation of genes involved in multiple categories including immune function, extracellular matrix, and cell proliferation in white adipose (**Figure 5 and Supplemental Table 4**).

**Figure 5.**
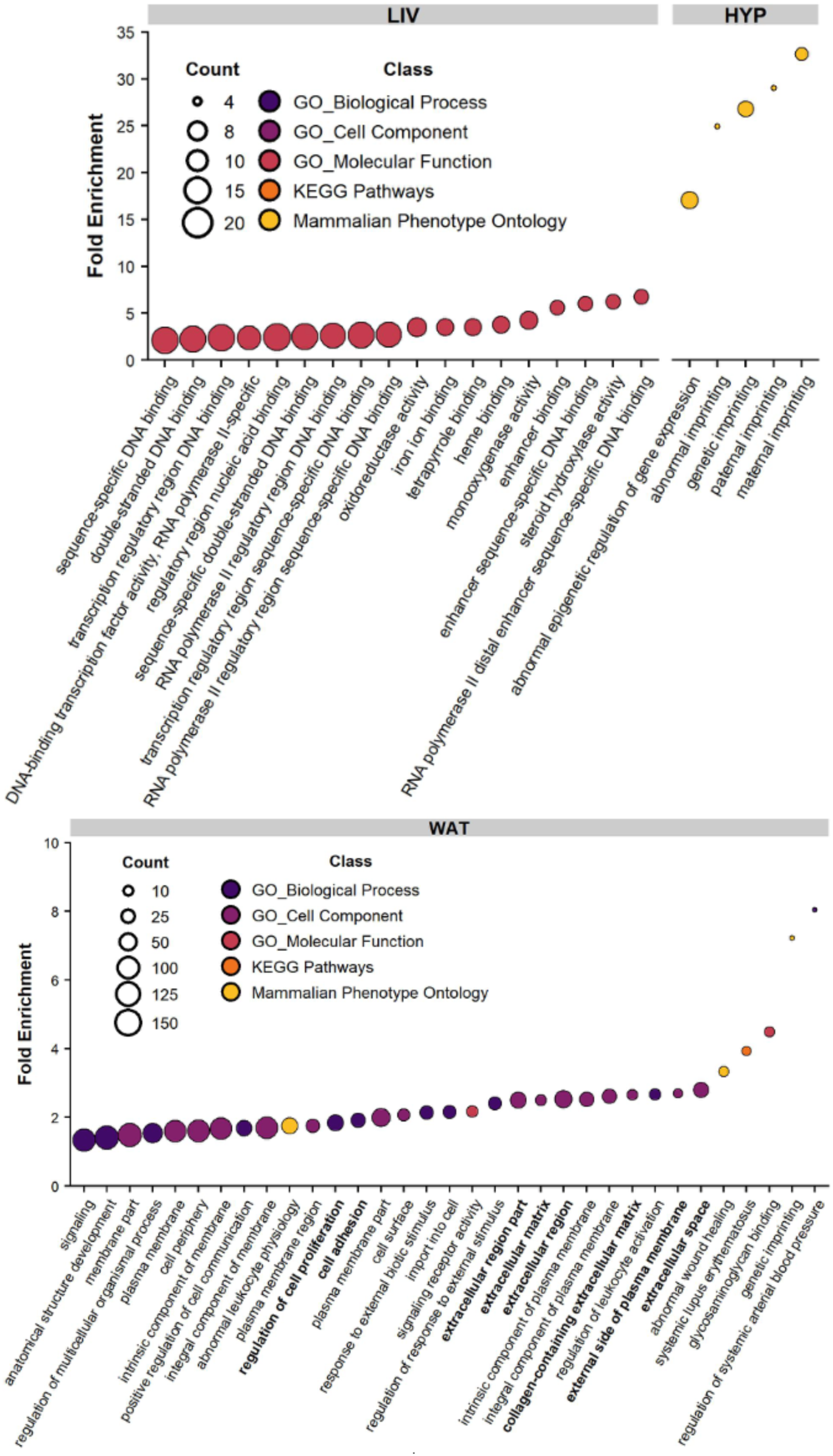
Enrichment analysis of interacting genes showing parent-of-origin effects. Over-representation of gene ontologies, KEGG pathways and Mammalian Phenotype ontologies in hypothalamus (HYP), white adipose (WAT) and liver (LIV).

### Epistasis in dietary-obesity QTL showing parent-of-origin effects

A subset of the genes in our significant interaction networks fall within the support intervals of our previously-identified metabolic QTL showing parent-of-origin effects on phenotype, so we tested for imprinting-by-imprinting epistasis in the F_16_ mapping data between interacting gene pairs where the gene showing biallelic expression fell within a metabolic QTL support interval and the gene showing parent-of-origin-dependent allele-specific expression had genotyped markers available. The numbers of unique biallelic genes that met this positional criteria for epistasis testing were 7 in hypothalamus, 45 in white adipose, and 11 in liver. The number of unique interacting genes showing parent-of-origin-dependent allele-specific expression with available markers were 28 in hypothalamus, 54 in white adipose, and 33 in liver (**Supplemental Table 5**).

In hypothalamus, we identified 9 significant epistatic interactions passing multiple tests correction among F_16_ genotyped markers that comprised 8 unique genes showing parent-of-origin-dependent allele-specific expression and 3 biallelic genes differentially expressed by cross (**Figure 6**). These genes were associated with three QTLs showing parent-of-origin effects: *Dserum17b*, associated with triglycerides; *Dserum1c*, associated with cholesterol; and *Ddiab8a*, associated with area under the curve for a glucose tolerance test (**Supplemental Table 5**).

**Figure 6.**
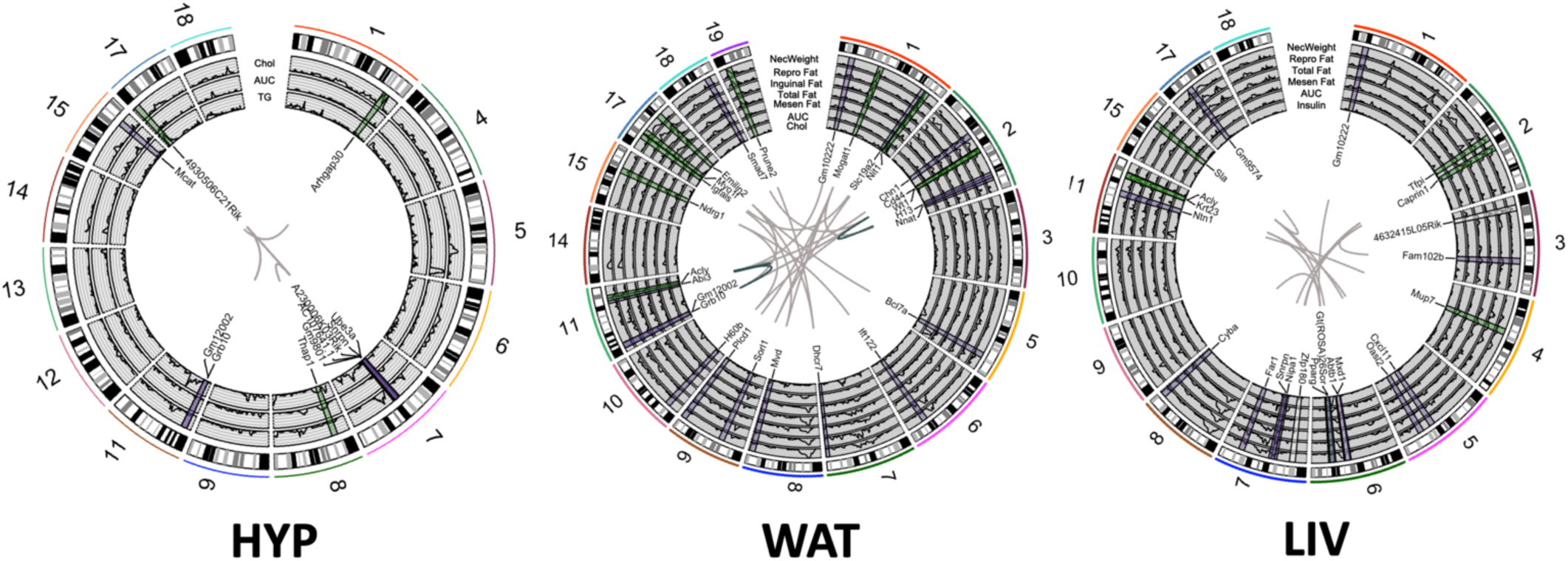
Networks of genes showing parent of origin effects show significant epistasis for dietary-obesity traits. Networks are comprised of two classes of parent-of-origin effect: parent-of-origin-dependent allele-specific expression (purple text and shading) and biallelic genes showing differential expression by cross, LxS or SxL (green text and shading). Inner tracks depict dietary-obesity QTL mapped in an F_16_ advanced intercross of the LG/J and SM/J inbred mouse strains. The outer track represents chromosomal position. Interacting gene pairs are connected by lines with *cis*-interactions colored purple and *trans*-interactions colored black. HYP = hypothalamus; WAT = white adipose; LIV = liver. AUC = area under the curve calculated from a glucose tolerance test; Chol = serum cholesterol; Inguinal Fat = inguinal fatpad weight; Mesen Fat = mesenteric fatpad weight; Nec Weight = weight at necropsy; Repro Fat = reproductive fatpad weight; Total Fat = total fatpad weights.

In white adipose, we identified 195 significant epistatic interactions passing multiple tests correction among F_16_ genotyped markers that comprised 16 unique genes showing parent-of-origin-dependent allele-specific expression and 10 biallelic genes differentially expressed by cross (**Figure 6**). These genes were associated with nine QTLs showing parent-of-origin effects: *Dob1b*, associated with inguinal, reproductive, and total fatpad weights and necropsy weight; *Dserum1c*, associated with cholesterol; *Dob2d*, associated with mesenteric and reproductive fatpad weights and necropsy weight; *Dob11b*, associated with reproductive and total fatpad weights and necropsy weight; *Dob15a*, associated with mesenteric fatpad weight; *Dserum17a*, associated with cholesterol; *Ddiab17b*, associated with area under the curve for a glucose tolerance test; *Dob17d*, associated with inguinal and reproductive fatpad weights; and *Dserum17a*, associated with cholesterol (**Supplemental Table 5**).

In liver, we identified 54 significant epistatic interactions passing multiple tests correction among F_16_ genotyped markers that comprised 12 unique genes showing parent-of-origin-dependent allele-specific expression and 7 biallelic genes differentially expressed by cross (**Figure 6**). These genes were associated with six QTLs showing parent-of-origin effects: *Dob2c*; associated with mesenteric fatpad weight; *Dob2d*, associated with reproductive fatpad weight and necropsy weight; *Ddiab4a*, associated with area under the curve for a glucose tolerance test; *Ddiab6d*, associated with serum insulin; *Dob11b*, associated with reproductive and total fatpad weights and necropsy weight; and *Dob15a*, associated with mesenteric fatpad weight (**Supplemental Table 5**).

Many of the parent-of-origin effects at these QTL are context dependent, and this is described in previous studies^41–44^.

### Interaction between Nnat and Mogat1 in white adipose tissue

*Nnat* (neuronatin) is a paternally-expressed imprinted gene encoding a lipoprotein involved in intracellular calcium signaling. In white adipose tissue, *Nnat* expression significantly covaries with *Mogat1* (p = 3.8e^-3^), a biallelic gene showing significant differential expression by cross in high fat-fed females. This differential expression pattern was validated by q-rtPCR in biological replicates (p = 0.01; **Supplemental Figure 2**). *Mogat1* (monoacylglycerol O-acyltransferase 1) is a gene encoding an enzyme that catalyzes the synthesis of diacylglycerol from monoacylglycerol, a precursor of triacylglycerol and other important lipids. *Mogat1* falls within the support intervals of the F_16_ QTL *Dob1b* which shows significant diet-by-sex parent-of-origin effects on multiple adiposity traits, including weight at necropsy and reproductive, inguinal, and total fatpad weights^41,44^. Genotyped markers around *Nnat* (UT-2-158.095429) and *Mogat1* (wu-rs13475931) show significant imprinting-by-imprinting epistasis for inguinal fatpad weight (p = 1.79e^-3^) and total fatpad weight (p = 2.5e^-4^) (**Figure 7**). Heterozygote F_16_ mice expressing the paternal LG/J-derived allele (SL) at *Nnat* have the highest total fatpad weight when they have the SL genotype at *Mogat1*. We find that NNAT and MOGAT1 expression are significantly correlated in human visceral adipose tissue (r=0.53, p<2.2e^-16^; **Supplemental Figure 3**), and MOGAT1 is an eQTL gene in human subcutaneous adipose tissue (rs140256046, p = 1.5e^-5^; GTEx)^46^.

**Figure 7.**
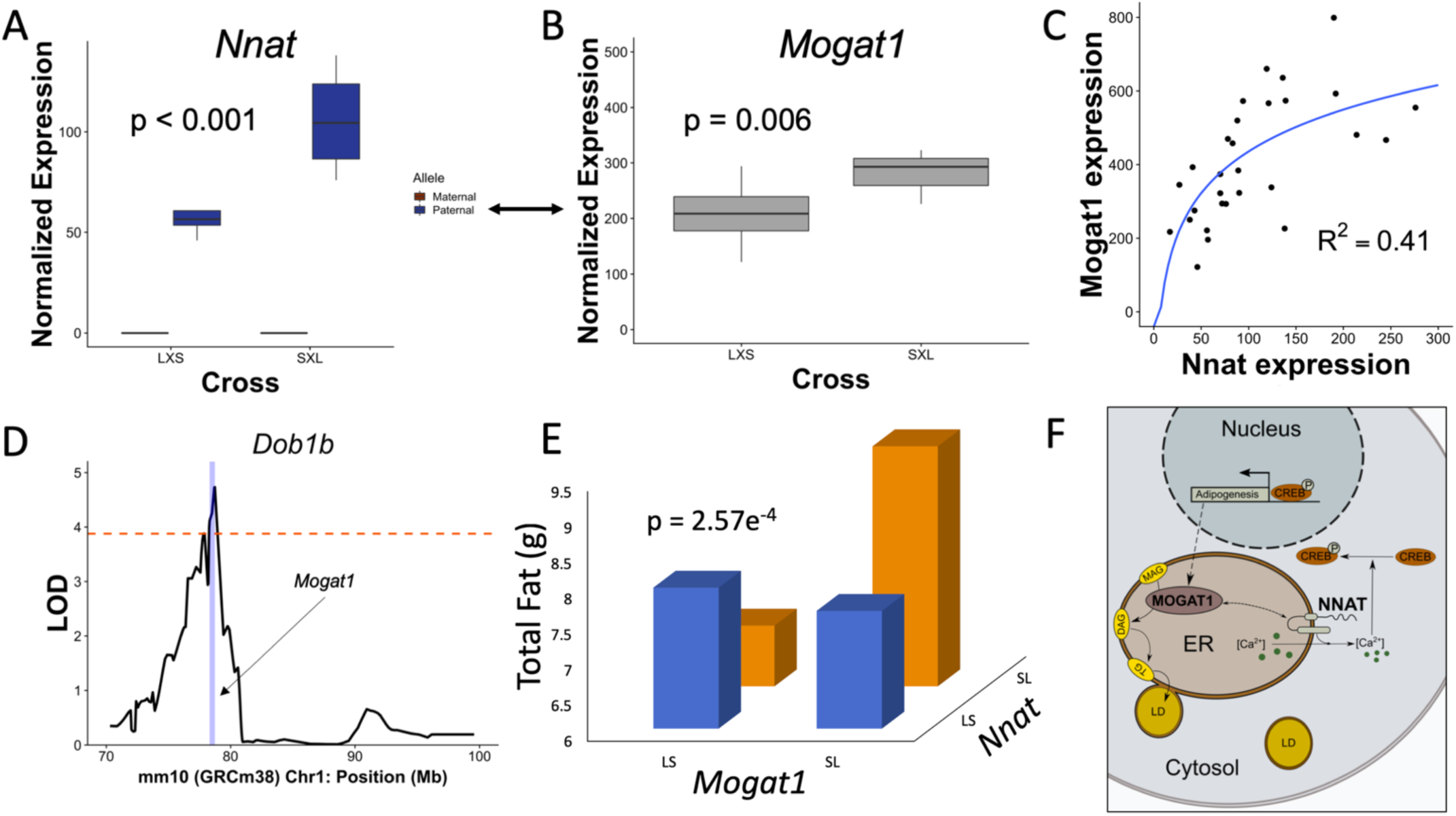
Interaction in white adipose between a gene showing parent-of-origin dependent allele-specific expression and a biallelic gene differentially expressed by cross. **A**. We identified *Nnat* as an imprinted gene showing paternal expression. **B**. *Mogat1* was identified as a biallelic gene showing differential expression between reciprocal crosses (LxS versus SxL), and **C**. found to significantly covary with *Nnat*. Logarithmic regression was performed fitting the function *Mogat1*∼ln(*Nnat*). **D**. *Mogat1* falls in a QTL for multiple adiposity traits identified in an F_16_ LGxSM advanced intercross. **E**. Targeted epistasis mapping in the F_16_ shows significant epistasis between *Nnat* and *Mogat1* for inguinal fatpad and total fat. Heterozygote mice expressing the paternal LG/J-derived allele (SL) at *Nnat* have the highest total fatpad weight when they have the SL genotype at *Mogat1*. **F**. *Nnat* and *Mogat1* both localize to the endoplasmic reticulum in adipocytes and contribute to adipose function.

## DISCUSSION

Epistatic interactions involving parent-of-origin effects on complex traits occur when the genotypic effects of one gene depends on the parent-of-origin of alleles at another gene^1^. In this study, we examined epistatic interactions associated with parent-of-origin effects on dietary-obesity traits in three tissues that are central to metabolism: hypothalamus, white adipose, and liver. We identified genes showing significant parent-of-origin-dependent allele-specific expression biases using a simple yet powerful F_1_ reciprocal cross mouse model. We found that they are highly context-specific, sometimes occurring in only one sex and/or diet cohort. Furthermore, the degree of context-specificity varies by tissue, with white adipose showing the highest degree of context-specific parent-of-origin effects on expression. This likely reflects adipose tissue’s intrinsic responsiveness to sex and its high degree of plasticity under different nutritional conditions^47–52^. Expression of canonically imprinted genes, such as *Peg10, Meg3, Grb10*, and *Snrpn*, show the lowest degree of context-specificity across all tissues. This likely reflects the fundamental role of these genes in growth and development^53–56^. Although these parent-of-origin dependent allele-specific expression biases are consistent with imprinting mechanisms, we cannot rule out that maternal and/or paternal effects also contribute to the phenomena we observe.

We identified biallelic genes that are significantly differentially expressed by cross (LxS or SxL) and found that they also show a high degree of context-specificity across all tissues. We then constructed interaction networks between these biallelic genes and those genes showing parent-of-origin-dependent allele-specific expression biases. We find that parent-of-origin effects are propagated through networks. Our proposed model is that genes showing parent-of-origin-dependent allele-specific expression biases affect the expression of biallelic genes differentially expressed by cross, which in turn contributes to parent-of-origin effects on phenotype. If environment affects the context-specificity of the parent-of-origin-dependent allele-specific expression biases, then the context-specificity would be propagated in the biallelic genes differentially expressed by cross. Indeed, we find that the subset of genes showing parent-of-origin-dependent allele-specific expression that significantly interact with biallelic genes that are differentially expressed by cross are enriched for those genes that are context-specific in white adipose (p = 0.001) and liver (p < 0.001). The proportion of context-specific genes showing parent-of-origin-dependent allele-specific expression biases increases from 41.18% overall to 48.8% of genes in networks in white adipose, and from 41.28% to 51.8% in liver. Hypothalamus, which has the lowest amount of context-specificity in genes showing parent-of-origin-dependent allele-specific expression biases, shows a non-significant decrease in context-specificity of interacting gene pairs (31.48% overall to 31.33%; p = 0.628).

Our results that genes showing parent-of-origin effects, both allele-specific expression biased and biallelic, form highly interconnected networks of covarying expression is consistent with previous studies showing that imprinted genes are particularly interactive^29,34,35^. We find that genes showing parent-of-origin-dependent allele-specific expression biases tend to cluster in genomic regions. This is expected of genes sharing *cis*-regulatory elements, as is the case in canonical imprinting, and is also consistent with other studies^33^. The degree of clustering varies among tissues; hypothalamus shows the highest degree of genomic clustering compared to white adipose and liver. Together with hypothalamus having the greater proportion of context-independent parent-of-origin-dependent allele-specific expression, we find that allele-specific expression patterns in the hypothalamus are most consistent with canonical imprinting. This is further validated by the enrichment analyses showing that interacting pairs of genes in hypothalamus are over-represented in categories related to imprinting (**Figure 5**).

Enrichment analyses in liver reveal interacting gene pairs that are over-represented in DNA binding and iron metabolism categories. Variation in iron metabolism is associated with variation in diabetes-related traits. Recently, the LG/J and SM/J inbred strains have been shown to vary in their glucose parameters in response to dietary iron^57–59^. Gene pairs in white adipose are over-represented in categories involved in extra-cellular matrix and cell proliferation. This is an intriguing finding because the extra-cellular matrix is integral for cellular signaling, regulation of growth factor bioavailability, and tissue remodeling^60,61^. The adipose extra-cellular matrix undergoes constant remodeling to accommodate changes in adipocyte shape and function in response to nutritional cues^62^. Disruption of adipose extra-cellular matrix components is associated with inflammation and tissue fibrosis as well as obesity-induced insulin resistance^63,64^. If genes showing parent-of-origin-dependent allele-specific expression act as intermediaries between environment (here, diet and/or sex) and cellular response, then interacting biallelic genes that are differentially expressed by cross may be a part of altered extra-cellular matrix or a product of that alteration. A previous study of interactions among imprinted genes in adipose also found an enrichment of genes involved in the extra-cellular matrix and cell proliferation^34^. Understanding the biological mechanisms and consequences of these interactions will improve our understanding of adipose biology and can direct research aimed at developing innovative therapeutic strategies for obesity-related metabolic disease.

We tested epistasis in an F_16_ advanced intercross of LG/J and SM/J mice (n=1002) between the significantly interacting gene pairs that we identified in our F_1_ LG/J x SM/J reciprocal cross. We only tested epistasis for gene pairs where the biallelic gene that is differentially expressed by cross fell within a dietary-obesity F_16_ QTL showing parent-of-origin effects on a metabolic phenotype. The previously reported QTL show a high degree of context-dependency of parent-of-origin effects, which is consistent with expression patterns of the F_1_ gene pairs we tested. We found more significant epistatic interactions between gene pairs identified in white adipose tissue than in either liver or hypothalamus. These interactions fall overwhelmingly in QTL associated with adiposity traits (instead of diabetic or lipid traits) and tend to be *trans*-interactions between genes on different chromosomes. This is consistent with the high proportion of *trans*-interactions seen in the overall interaction networks for white adipose tissue between genes showing parent-of-origin-dependent allele-specific expression and biallelic genes that are differentially expressed by cross. In some cases, the epistasis is between marker pairs showing linkage disequilibrium, suggesting that the parent-of-origin effects at some QTL is propagated through linkage between interacting genes. Significant interacting pairs where linkage disequilibrium is not detected may represent novel QTL that previous studies were underpowered to detect.

Many of the genes we identify may interact on a cellular or physiological level to affect dietary-obesity traits. Of the 28 epistatic network members identified in white adipose, only 8 have no obvious connection to adiposity traits (*Gm10222, Gm12002, Nit1, Ift122, Abi3, H60b, Prune2*, and *Myo1f*). Others are involved in adipocyte differentiation (*Mogat1, Bcl7a, Ndrg1, Emilin2, Nnat*, and *Smad7*), variation in obesity, insulin, and glucose traits (*Slc19a2, Chn1, Wt1, H13, Dhcr7, Cd44, Grb10*, and *Igfals*), and/or play important functions in the synthesis and degradation of lipids (*Plcd1, Mogat1, Mvd, Acly, Sorl1*, and *Dhcr7*). Of particular interest is the interaction between the paternally imprinted gene *Nnat* (neuronatin) and *Mogat1* (monoacylglycerol O-acyltransferase 1).

*Mogat1* falls in a QTL (*Dob1b*) that shows significant diet-by-sex parent-of-origin effects on multiple adiposity traits: reproductive and inguinal fatpad weights, total fatpad weight, and weight at necropsy^41,44^. Both *Nnat* and *Mogat1* localize in the endoplasmic reticulum (ER) of white adipose tissue. *Nnat* is a diet-responsive proteolipid known to play a role in ER calcium efflux as a part of Ca^2+^ signaling^65^. Increased cytosolic Ca^2+^ levels lead to CREB (cAMP response element-binding protein) activation, which in turn promotes adipogenesis. *Mogat1* expression is induced during adipogenesis. *Mogat1* plays an important role in adipocyte differentiation by contributing to the production and accumulation of triglycerides by catalyzing the conversion of mono-acylglycerides to di-acylglycerides, which are subsequently converted to tri-acylglycerides^66^. The correlation between Nnat and Mogat1 is non-linear, which supports our proposed model that *Nnat* affects *Mogat1* by regulating activation of CREB. In this scenario, CREB expression is a limiting step in the transduction of signal from *Nnat* to *Mogat1*. At some point increased *Nnat* expression would cease to increase *Mogat1* expression because all available CREB has been activated. This would appear as the plateau we observe in *Mogat1* expression (**Figure 7C**). MOGAT1 is an eQTL in human subcutaneous adipose tissue, which is homologous to the mouse inguinal fatpad^46^. Further, NNAT and MOGAT1 expression are significantly correlated in human visceral adipose tissue, with a pattern similar to that observed in mice (**Supplemental Figure 3**). Thus, there is good evidence that both of these genes contribute to adiposity, that they may interact indirectly through CREB activity, and that this interaction contributes to the parent-of-origin effects on adiposity observed in the F_16_ QTL.

The support intervals for the original F_16_ parent-of-origin QTL span large genomic regions and are not empowered to identify the specific genes that contribute to the parent-of-origin-dependent phenotypic variation. By leveraging the reciprocal F_1_ hybrids, we are able to integrate parent-of-origin-dependent allele-specific expression and parent-of-origin-dependent differential expression with the mapped F_16_ phenotypes. By further incorporating multiple metabolically-relevant tissues as well as sex and dietary environments, we provide an authentic systems biology perspective on metabolic trait variation. By doing so, we identify plausible candidates for functional validation and describe discrete molecular networks that may contribute to the observed parent-of-origin effects on metabolic traits. The genes and networks we present here represent a set of actionable interacting candidates that can be probed to further identify the machinery driving these phenomena and make predictions informed by genomic sequence. The frameworks we have developed account for the genetic, epigenetic, and environmental components underlying these parent-of-origin effects, thereby improving our ability to predict complex phenotypes from genomic sequence. We focused on metabolic phenotypes in this study, but the patterns we identified may translate to other complex traits where parent-of-origin effects have been implicated.

## METHODS

### Mouse husbandry and phenotyping

LG/J and SM/J founders were obtained from The Jackson Laboratory (Bar Harbor, ME). F_1_ reciprocal cross animals were generated by mating LG/J mothers with SM/J fathers (LxS) and the inverse (SxL). At three weeks of age, animals were weaned into same-sex cages and randomly placed on high-fat (42% kcal from fat; Teklad TD88137) or low-fat (15% kcal from fat; Research Diets D12284) isocaloric diets. Animals were weighed weekly until sacrifice. At 19 weeks of age, body composition was determined by MRI and a glucose tolerance test was performed after a 4 hour fast. At 20 weeks of age, animals were given an overdose of sodium pentobarbital after a 4 hour fast and blood was collected via cardiac puncture. Euthanasia was achieved by cardiac perfusion with phosphate-buffered saline. After cardiac perfusion, liver, reproductive fatpad and hypothalamus were harvested, flash frozen in liquid nitrogen, and stored at −80°C.

### Genomes and annotations

LG/J and SM/J indels and SNVs were leveraged to construct strain-specific genomes using the GRC38.72-mm10 reference as a template^67^. This was done by replacing reference bases with alternative (LG/J | SM/J) bases using custom python scripts. Ensembl R72 annotations were adjusted for indel-induced indexing differences for both genomes.

### RNA sequencing

Total RNA was isolated from adipose and hypothalamus tissues using the RNeasy Lipid Tissue Kit (QIAgen) and from liver using TRIzol (n = 32, 4 animals per sex/diet/cross cohort). RNA concentration was measured via NanoDrop and RNA quality/integrity was assessed with a BioAnalyzer (Agilent). RNA-Seq libraries were constructed using the RiboZero kit (Illumina) from total RNA samples with RIN scores >8.0. Libraries were checked for quality and concentration using the DNA 1000LabChip assay (Agilent) and quantitative PCR, according to manufacturer’s protocol. Libraries were sequenced at 2×100 paired end reads on an Illumina HiSeq 4000. After sequencing, reads were de-multiplexed and assigned to individual samples.

### Allele-specific expression

FASTQ files were filtered to remove low quality reads and aligned against both LG/J and SM/J pseudo-genomes simultaneously using STAR with multimapping disallowed^68^. Read counts were normalized via upper quartile normalization and a minimum normalized read depth of 20 was required. Alignment summaries are provided in **Supplemental Table 6 and Supplemental Figure 2**.

For each gene in each individual, allelic bias (L_bias_) was calculated as the proportion of total reads for a given gene with the LG/J haplotype. Parent-of-origin-dependent allele-specific expression was detected by ANOVA using one of a number of models in which L_bias_ is responsive to cross and some combination of sex and diet:

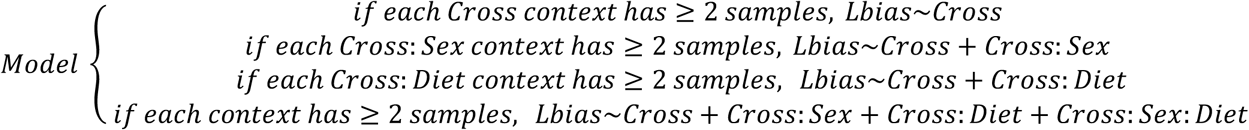

Accurately estimating the significance of these effects is challenging for two reasons: 1) the complexity of the many environmental contexts, and 2) the correlation of allelic bias within and between imprinted domains breaks assumptions of independence. A permutation approach is an effective way to overcome these challenges. The context data was randomly shuffled and analyses were rerun in order to generate a stable null distribution of F-statistics (**Supplemental Figure 3**). Significance thresholds were calculated from the empirical distribution of this null model and a p-value ≤ 0.05 was considered significant (**Supplemental Table 1**).

To determine the parental direction and size of expression biases, a parent-of-origin effect POE score was calculated as the difference in mean L_bias_ between reciprocal crosses (LxS or SxL). POE scores range from completely maternally-expressed (−1), to biallelic (0), to completely paternally-expressed (+1). POE score thresholds were calculated from a critical value of α = 0.01, determined from a null distribution created by permutation (**Supplemental Figure 4**). Genes with significant allele-specific expression and parent-of-origin scores beyond the critical value were considered to have significant parent-of-origin-dependent allele-specific expression.

### Library complexity

Complexity was measured by fitting a beta-binomial distribution to the distribution of L_bias_ values using the VGAM package^69^. The shape parameters (α, β) of beta-binomial distributions were estimated and used to calculate dispersion (ρ). Dispersion values less than 0.05 indicate our libraries are sufficiently complex (**Supplemental Figure 5**).

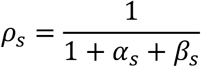

Two libraries, one from white adipose and one from liver, were found to have insufficient complexity and were removed from the analyses.

### Differential expression

Differential expression by reciprocal cross was determined by first aligning reads against the LG/J and SM/J genomes simultaneously with multimapping permitted. Reads were normalized by TMM and a minimum normalized read count of 10 was required. Generalized linear models accounting for diet, sex, and diet-by-sex were fit in EdgeR^70^. Differential expression was detected by a likelihood ratio test. Significance was determined for four nested models for each gene:

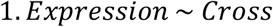

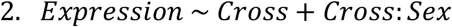

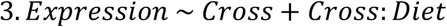

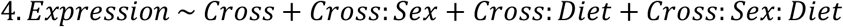

To accurately estimate significance given the complexity of the many diet and sex cohorts, the context data was shuffled and the analyses rerun in order to generate an appropriate null distribution of likelihood ratio statistics for each model (**Supplemental Figure 6**). Stability of the permuted data was evaluated by calculating the total quantile deviation (TQD) at each iteration:

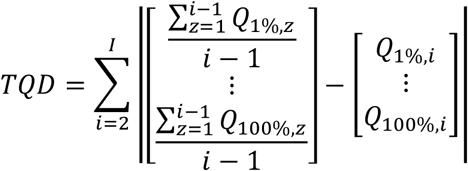

Genes with a p-value of ≤ 0.05 and a |*log*_2_(*fold change*)| ≥ 1 were considered significantly differentially expressed by reciprocal cross (**Supplemental Figure 7 and Supplemental Table 2**).

### Gene-gene interactions

Networks were constructed in each tissue by pairing genes showing parent-of-origin-dependent allele-specific expression with biallelic genes that are differentially expressed by cross. Pairs were predicted by modeling the expression of biallelic genes as a function of parent-of-origin-dependent allele-specific expression, L_bias_, sex, diet, and sex-by-diet. The strength of a prediction was measured through model fit, which was estimated as a mean test error with 10-fold cross-validation employed to prevent overfitting. Given the complexity of contexts, the significance threshold was determined by permuting the context data to generate a stable null-distribution of mean test errors (**Supplemental Figure 8**). A p-value ≤ 0.01 was considered significant (**Supplemental Table 3**). Genomic clusters of interacting genes were assigned by K-means clustering performed for each chromosome separately.

### Functional enrichment analysis

Functional enrichment of interacting genes showing parent-of-origin-dependent allele-specific expression with biallelic genes that are differentially expressed by cross was tested by over-representation analysis in the WEB-based Gene Set Analysis Toolkit v2019^71^. We performed analyses of gene ontologies (biological process, cellular component, molecular function), pathway (KEGG), and phenotype (Mammalian Phenotype Ontology). For each tissue, the list of all unique interacting genes was analyzed against the background of all unique genes expressed in that tissue. A Benjamini-Hochberg FDR-corrected p-value ≤ 0.01 was considered significant (**Supplemental Table 4**).

### Enrichment in QTL with genomic imprinting effects

Support intervals of quantitative trait loci (QTL) showing significant genomic imprinting effects were randomly shuffled throughout the genome to generate an empirical distribution of random QTL-sized regions. The following gene sets were intersected with these random regions and compared to their intersections in the mapped QTL: i) known imprinted genes; ii) genes differentially expressed by reciprocal cross; iii) genes showing parent-of-origin-dependent allele-specific expression; and iv) gene-gene interactions between genes differentially expressed by reciprocal cross and genes showing parent-of-origin-dependent allele-specific expression (**Supplemental Tables 1, 2, 3, and 7**). P-values ≤ 0.05 were considered significant and ≤ 0.1 considered suggestive.

### Epistasis testing

The F_16_ LxS advanced intercross population, phenotypes, genotypes, genotypic scores, and QTL mapping methods are described elsewhere^41–44^. We tested for epistasis in interacting pairs between genes showing parent-of-origin-dependent allele-specific expression and biallelic genes that are differentially expressed by cross where the biallelic gene falls within a QTL showing parent-of-origin effects. We selected F_16_ genotyped markers that fall within 1.5mB up-and downstream from the geometric center of each gene, defined as the genomic position halfway between the transcription start and stop position of that gene (**Supplemental Table 5, Supplemental Table 8**). For every F_16_ animal, an “imprinting score” was assigned to each marker based on that animal’s genotypic values (LL = 0, LS = 1, SL = -1, SS = 0; maternal allele is depicted first). Non-normally distributed phenotypes (as evaluated by a Shapiro-Wilk test) were log_10_-transformed to approximate normality. Because of the number of epistasis tests performed and the number of contexts represented in the data, we removed the effects of sex, diet and their interaction from each F_16_ phenotype with a covariate screen. We tested for epistasis on the residualized data using the following generalized linear model:

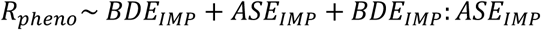

Where *R*_*pheno*_ is the residual phenotype, *BDE*_*IMP*_ is the imprinted genotypic score for the biallelic gene that is differentially expressed by cross, *ASE*_*IMP*_ is the imprinted genotypic score for the gene showing parent-of-origin-dependent allele-specific expression bias, and *BDE*_*IMP*_*:ASE*_*IMP*_ is the interaction between the two genes’ imprinted genotypic score. We employed a permutation approach in order to accurately estimate significance given the high correlation between metabolic phenotypes and the linkage of proximal markers. Imprinted genotypic values were randomly shuffled to generate a stable null model of F-statistics. Significance was calculated from the empirical cumulative distribution of the null model (**Supplemental Figure 9**). A p-value < 0.05 was considered significant. Epistasis was considered significant if the *BDE*_*IMP*_: *ASE*_*IMP*_ interaction term met the significance threshold. For all pairs showing significant epistasis, linkage disequilibrium was tested using Pegas^72^ (**Supplemental Table 5**).

### Validation of Nnat and Mogat1 differential expression

Differential expression of *Nnat* in white adipose was validated by q-rtPCR in LG/J and SM/J mice and in biological replicates of F_1_ reciprocal cross animals (n = 6 LG/J homozygotes, n = 8 LxS and 7 SxL reciprocal heterozygotes, n = 6 SM/J homozygotes). Differential expression in white adipose of the biallelic gene Mogat1 was also validated in reciprocal heterozygotes (n = 8 LxS and 7 SxL; **Supplemental Figure 2**). Total RNA was extracted from adipose samples using the Qiagen RNeasy Lipid Kit. High-Capacity cDNA Reverse Transcription Kit (Thermofisher) was used for reverse transcription. Quantitative-rtPCR was performed with an Applied Biosystems (USA) QuantStudio 6 Flex instrument using SYBR Green reagent. Results were normalized to L32 expression using the ΔΔC_t_ method. *Nnat* forward primer – CTACCCCAAGAGCTCCCTTT and reverse primer – CAGCTTCTGCAGGGAGTACC. *Mogat1* forward primer – TGTCTTGTCAAAACGCAGGAT and reverse primer – ACAACGGGAAACAGAACCAGA. L32 forward primer – TCCACAATGTCAAGGAGCTG and reverse primer – GGGATTGGTGACTCTGATGG. Data points were considered outliers if they led to violation of normality assumptions or were considered outliers by box and whisker plots. Significance was estimated using the students t-test.

### Correlation of NNAT and MOGAT1 expression in human adipose

Human visceral adipose expression raw count data was downloaded from the GTEx website on April 22, 2019^46^. Expression was normalized by TMM as implemented in EdgeR^70^. Normalized expression was log transformed and co-expression between NNAT and MOGAT1 was determined by Pearson’s product-moment correlation, n = 340 individuals (**Supplemental Figure 3**). Logarithmic regression was performed by fitting the function MOGAT1∼ln(NNAT).

## DATA ACCESS

All data generated and/or analyzed during the current study are available in the Supplemental Materials, in referenced publications, and at lawsonlab.wustl.edu. RNAseq reads generated for this study have been submitted to the NCBI Gene Expression Omnibus. Code written to analyze data is available on GitHub (https://github.com/LawsonLab-WUSM/POE_Epistasis)

## ACKNOWLEDGMENTS

This work was supported by NIDDK-K01DK095004, NIDDK-P30DK056341, NIDDK-P30DK020579, and NHGRI-T32-GM007067.

## AUTHOR CONTRIBUTIONS

HAL and JFM conceived of the study. HAL and JPW generated and collected data. LY, LS, and KF assisted in tissue harvesting. MAM performed q-rtPCR. HAL, JFM, and CLS performed analyses. JMC and CFS provided resources. HAL, JFM, and CLS wrote the manuscript. All authors read and approved the submitted manuscript.

## Supplemental Materials

**Supplemental Figure 1:** Metabolic QTL showing parent of origin effects are not enriched for imprinted genes.

**Supplemental Figure 2:** Validation of *Nnat* and *Mogat1* differential expression in biological replicates.

**Supplemental Figure 3:** Correlated expression of NNAT and MOGAT1 human visceral adipose.

**Supplemental Figure 4:** Number of reads mapped to LG/J x SM/J pseudo-genome. **Supplemental Figure 5:** Stable null permutation plots for allele-specific expression. **Supplemental Figure 6:** Permutation plots for parent-of-origin effect scores.

**Supplemental Figure 7:** RNAseq libraries are sufficiently complex to detect allele specific expression.

**Supplemental Figure 8:** Stable null permutation plots for differential expression by cross.

**Supplemental Figure 9:** Volcano plots of differentially expressed genes

**Supplemental Figure 10:** Stable null permutation plots for network pairs.

**Supplemental Figure 11:** Stable null permutations plot for epistasis.

**SupplementalTable1.xlsx:** Allele-specific expression

**SupplementalTable2.xlsx:** Biallelic genes differentially expressed by cross **SupplementalTable3.xlsx:** Networks of genes showing parent-of-origin allele-specific expression interacting with biallelic genes that are differentially expressed by cross

**SupplementalTable4.xlsx:** Over-representation input/output

**SupplementalTable5.xlsx:** Epistasis results

**SupplementalTable6.xlsx:** Alignment summaries

**SupplementalTable7.xlsx:** List of imprinted genes queried

**SupplementalTable8.xlsx:** QTL traits, positions, and genes falling within support intervals

