## Supplemental Figures for "Epistatic Networks Associated with Parent-of-Origin Effects on Metabolic Traits"

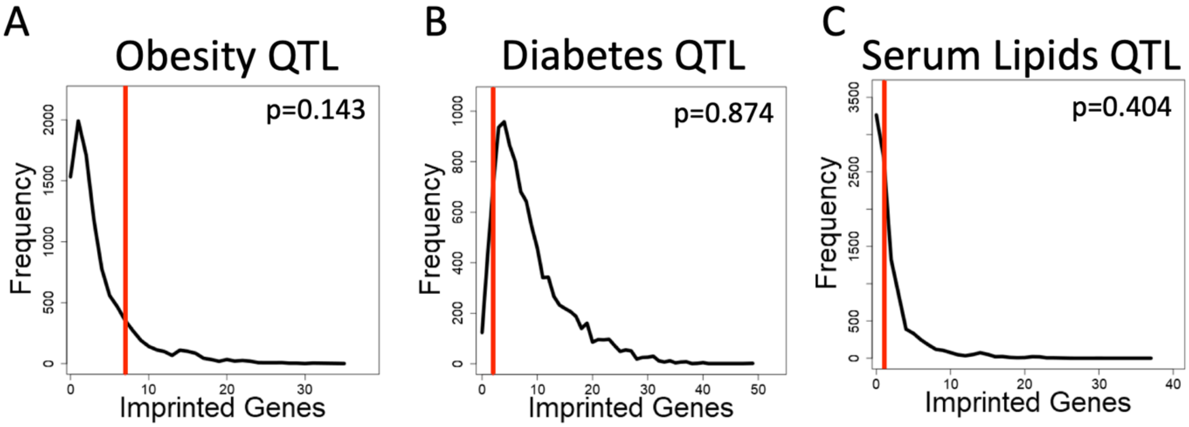


**Supplemental Figure 1: Metabolic QTL showing parent of origin effects are not enriched for imprinted genes**. Permutation analysis was performed to test for enrichment of canonically imprinted genes in metabolic QTL showing parent of origin effects. All obesity (**A**), diabetes (**B**), and serum lipids (**C**) QTL showing parent-of-origin effects were considered. QTL position was shuffled 10,000 times to establish a null model of imprinted genes within genomic regions of QTL interval sizes. The true number of canonically imprinted genes intersecting QTL showing a parent of origin effect is denoted with a vertical red line.

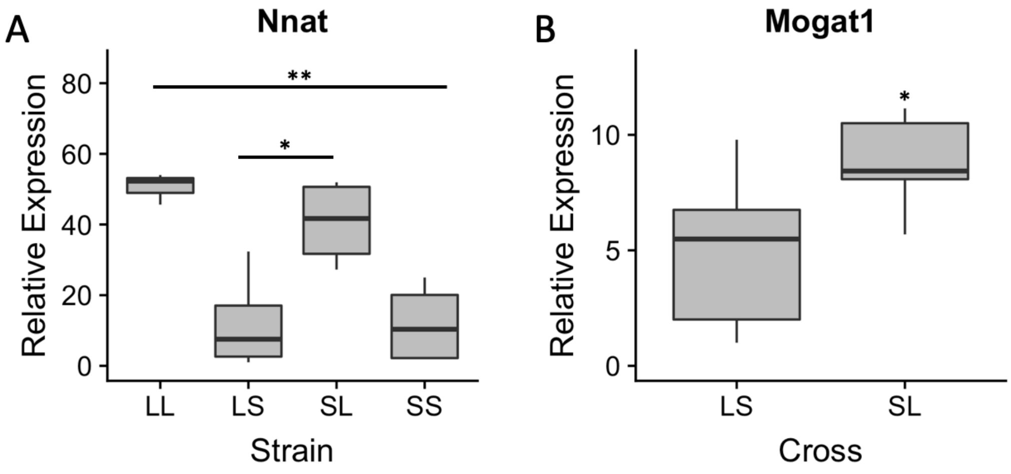


**Supplemental Figure 2: Validation of *Nnat* and *Mogat1* differential expression in biological replicates.** Differential expression in white adipose (reproductive fatpad) of the paternally expressed *Nnat* (**A**) and biallelic gene *Mogat1* (**B**) was validated by q-rtPCR in biological replicates (n = 6 LG/J homozygotes, n = 8 LxS and 7 SxL reciprocal heterozygotes, n = 6 SM/J homozygotes; p = 0.01).

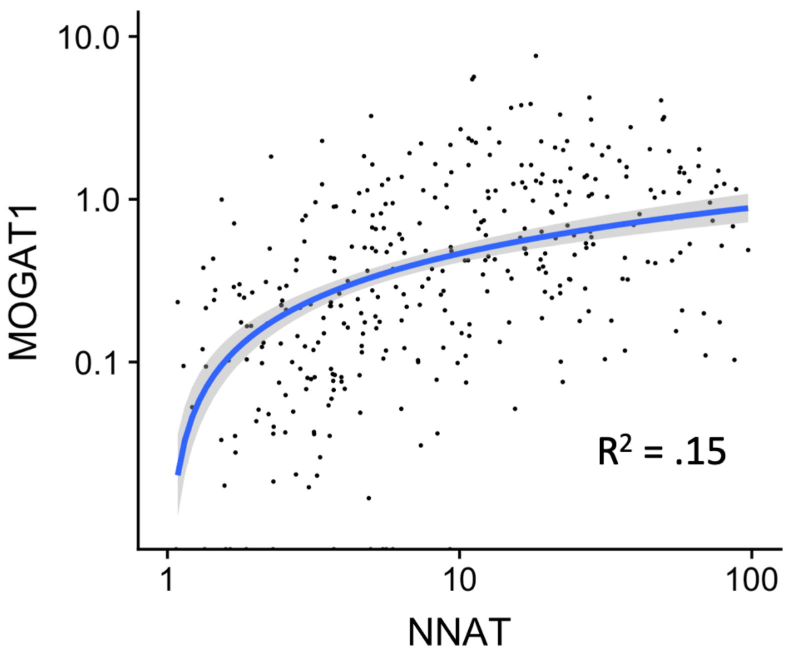


**Supplemental Figure 3: Correlated expression of NNAT and MOGAT1 human visceral adipose.** MOGAT1 and NNAT are significantly co-expressed in human omental adipose tissue, p <2.2e$^{-16}$, n = 340. Data from GTEx (https://gtexportal.org/home/).

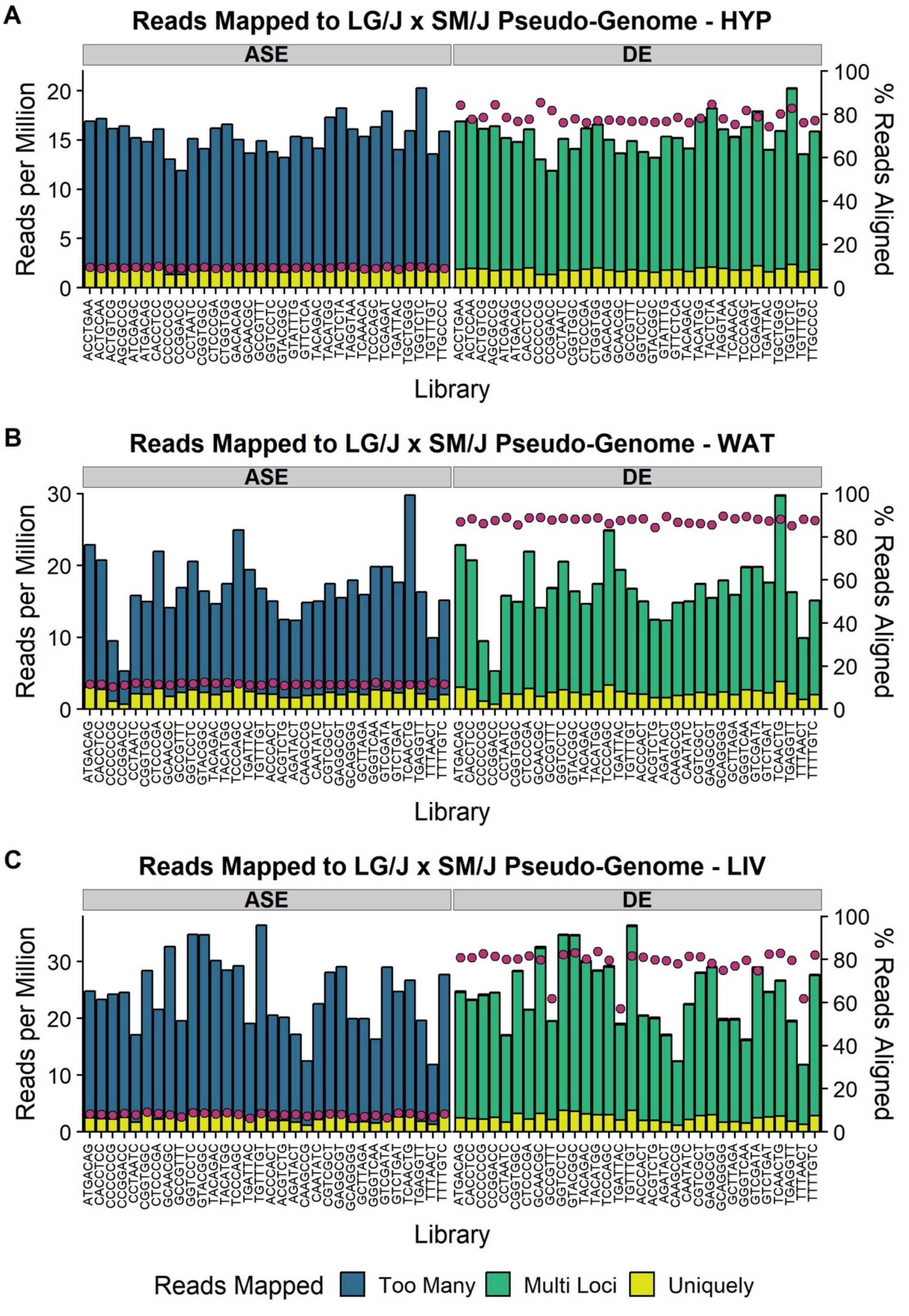


**Supplemental Figure 4: Number of reads mapped to LG/J x SM/J pseudo-genome.** We isolated total RNA from hypothalamus, white adipose, and liver (n = 32 per sex/diet/cross cohort) and constructed RNA-Seq libraries. Libraries were sequenced 1 sample per lane on an Illumina HiSeq 4000 (Illumina, San Diego, USA) using 2x100bp PE reads. FASTQ files were filtered to remove low quality reads. Remaining reads were aligned against both LG/J and SM/J pseudo-genomes simultaneously using STAR with either multimapping disallowed (allele-specific expression, **ASE**) or multimapping allowed (differential expression, **DE**). Libraries for each tissue are plotted separately: (**A**) hypothalamus, (**B**) white adipose, and (**C**) liver. The stacked bars denotes the number of reads per million (left y-axis) and are color-coded by category (uniquely = yellow, multi-loci = green, too many = blue). For the ASE alignment, “too many” reads are defined as n > 1 since multimapping was disallowed. For the DE alignment, “multi-loci” reads are defined as 1 < n < 10 and “too many” reads are defined as n > 10. The pink dots denote the primary alignment rates (right y-axis) for each library. Alignment rates are defined as (# reads mapped uniquely + # reads mapped multi-loci / total # input reads).

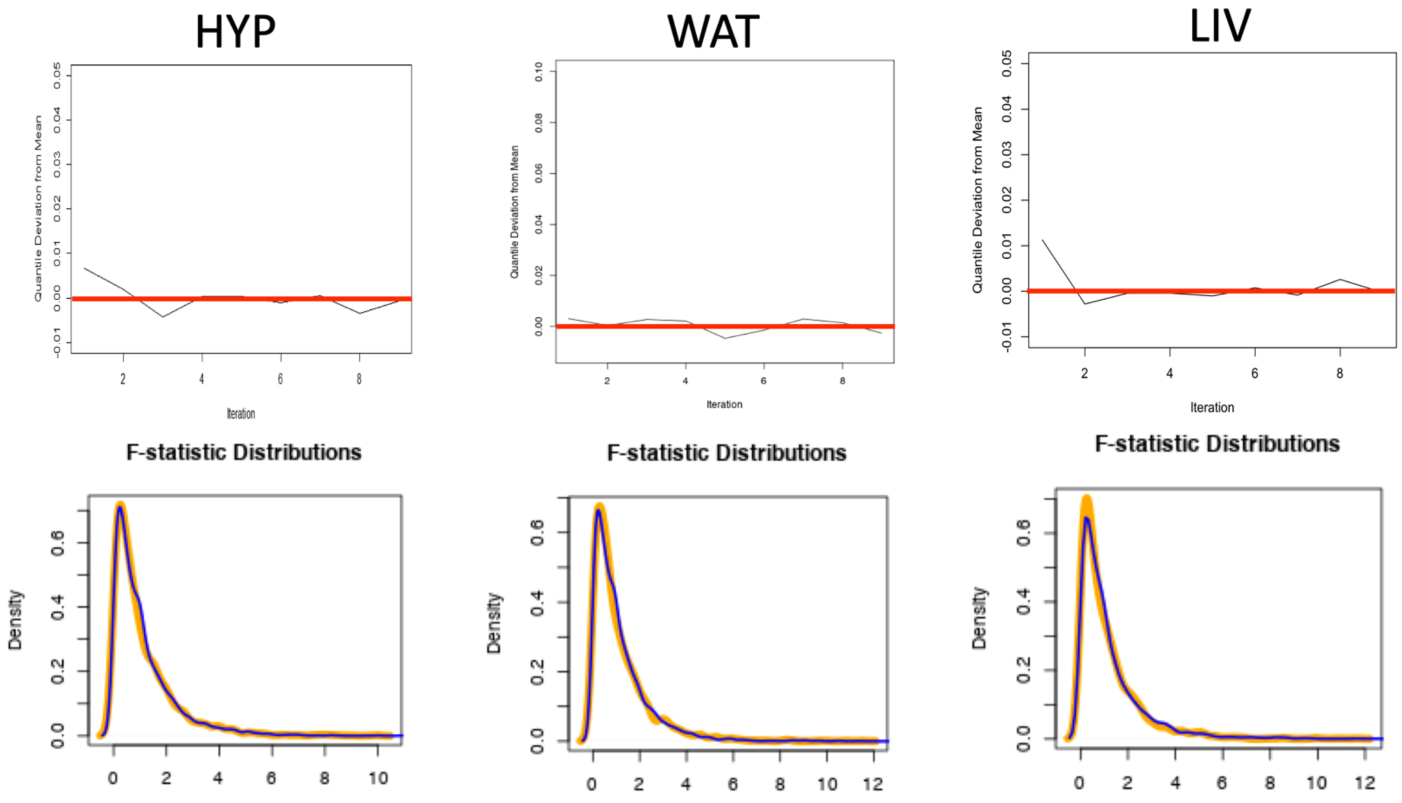


**Supplemental Figure 5: Stable null permutation plots for allele-specific expression.** Context was permuted to generate an appropriate null model of F-statistics. Null distribution stability was evaluated by calculating the total quantile deviation (TQD) at each iteration. TQD is the sum of deviations of quantile values (Q) from the mean of quantile values of all prior iterations. Quantiles corresponded to one percent increments of the distribution. The TQD of F-statistics under the null model approaches zero (red line) over ten iterations. This approximation towards zero demonstrates the stabilization of the null model. By ten shuffles the TQD value fluctuates ~0.005 units per iteration in the most complex model, meaning that at worst F-statistics estimates vary by ± 0.005, which corresponds to a p-value estimate error of ± $3.57 \times 10^{-5}$ applying the standard F-distribution to estimate significance. The null distribution of F-statistics (**Orange**) is plotted against the observed F-statistic distribution (**Blue**) for all tissues. Hyp = hypothalamus; WAT = white adipose; LIV = liver.

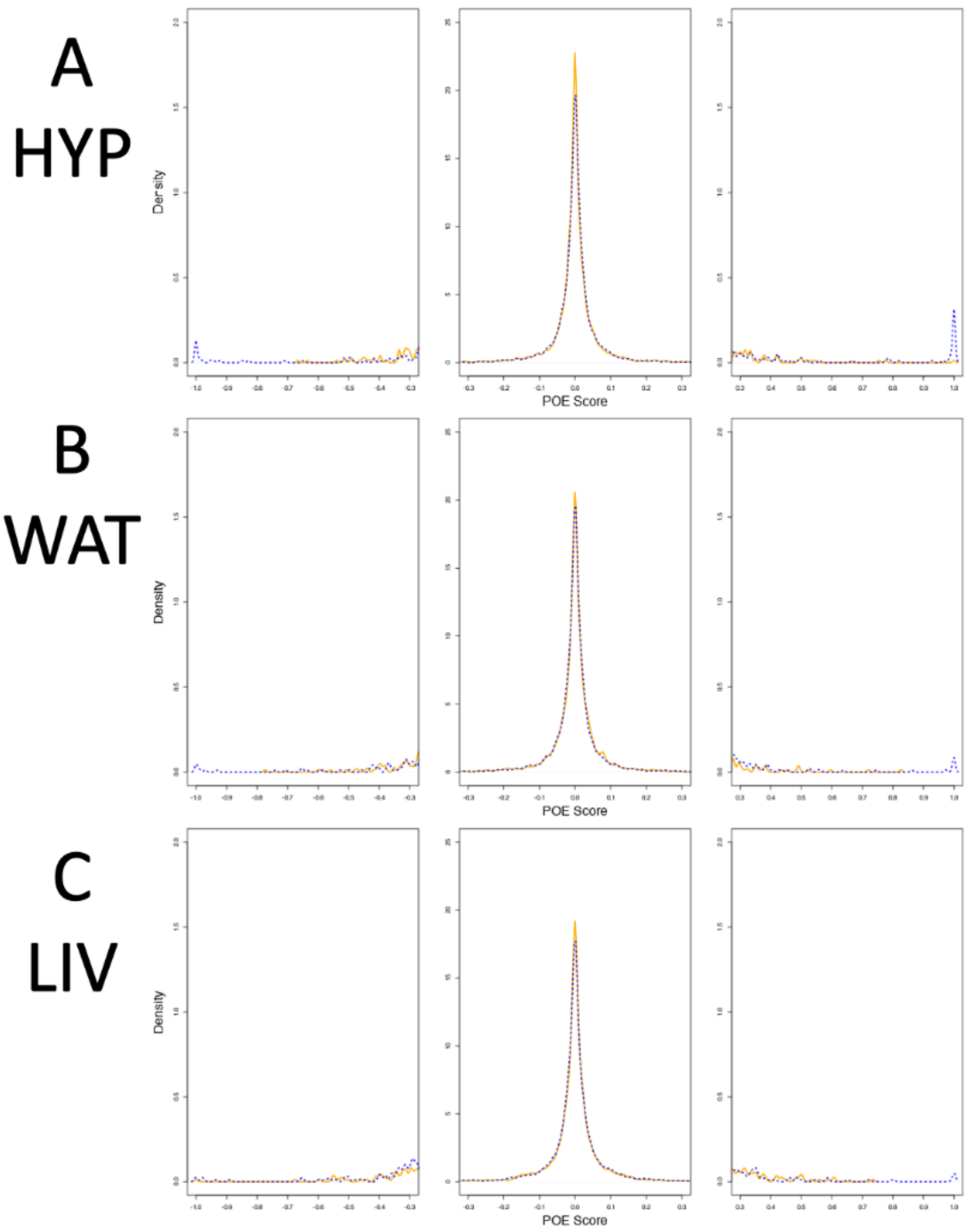


**Supplemental Figure 6: Permutation plots for parent-of-origin effect scores.** The parental direction and magnitude of expression bias is measured as a parent-of-origin effect (POE) score. POE scores were calculated as the difference in mean $L_{bias}$ between reciprocal crosses (LxS or SxL). POE scores range from completely maternally-expressed (-1), to biallelic (0), to completely paternally-expressed (+1). This was performed in hypothalamus (**A**), white adipose (**B**), and liver (**C**). The distribution under a null model of POE scores was generated by shuffling contexts. As expected, the null distribution of POE scores (orange) has more low values closer to zero than the real data distribution (blue). Thresholds were calculated from a critical value of α = 0.01, determined from the null distribution of POE scores for each tissue. Hyp = hypothalamus; WAT = white adipose; LIV = liver.

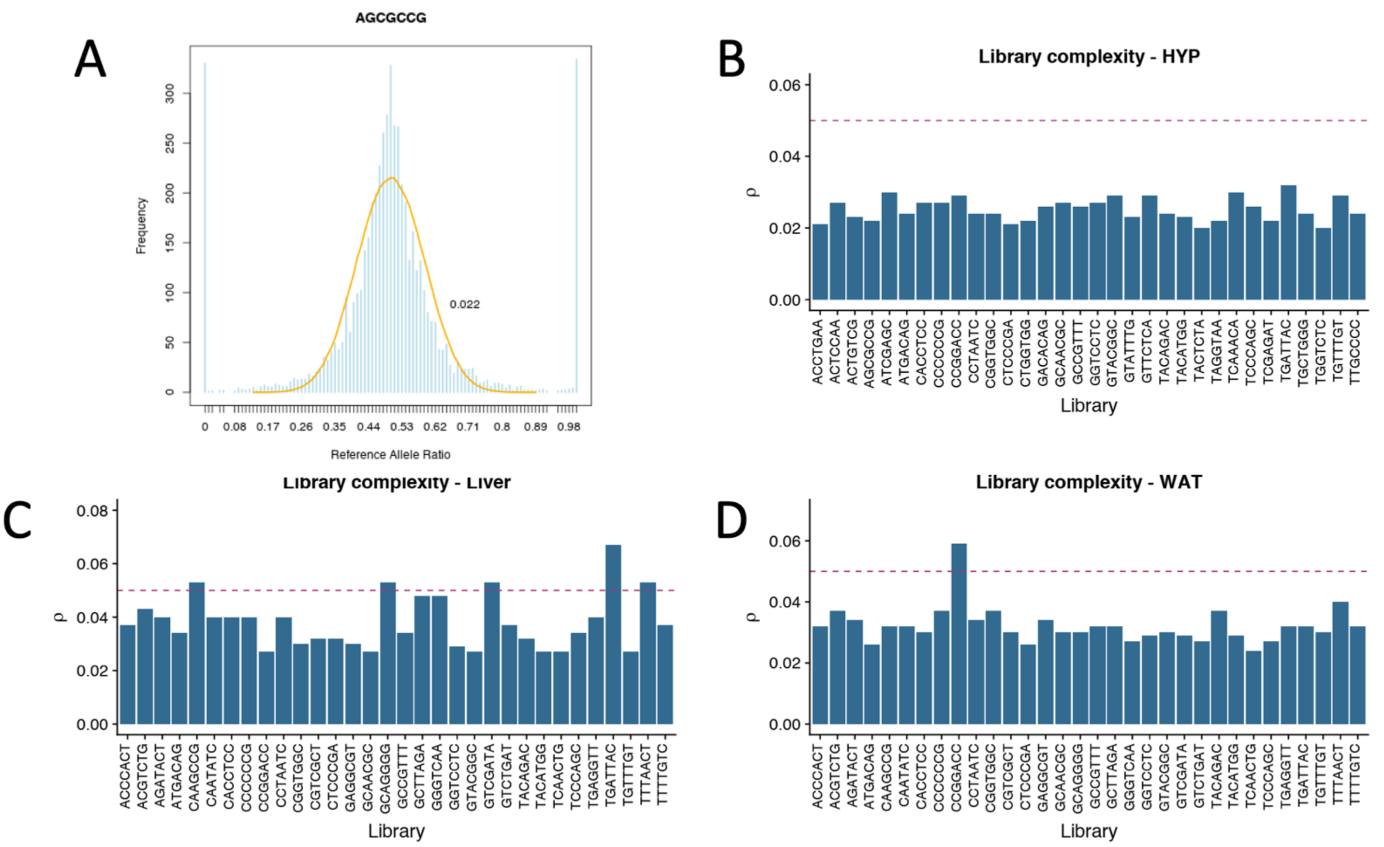


**Supplemental Figure 7: RNAseq libraries are sufficiently complex to detect allele specific expression.** Complexity was measured by fitting a beta-binomial distribution to the distribution of $L_{bias}$ values. A representative distribution (**A**) of $L_{bias}$ values (blue) and fit beta-binomial (orange). The shape parameters (α, β) of the beta-binomial distribution are used to calculate dispersion (ρ). Dispersion value less than 0.05 indicate sufficient complexity. All hypothalamus libraries were sufficiently complex (**B**). Two libraries were deemed insufficiently complex, one from Liver (**C**) one from white adipose (**D**), and were removed from analyses. Hyp = hypothalamus; WAT = white adipose; LIV = liver.

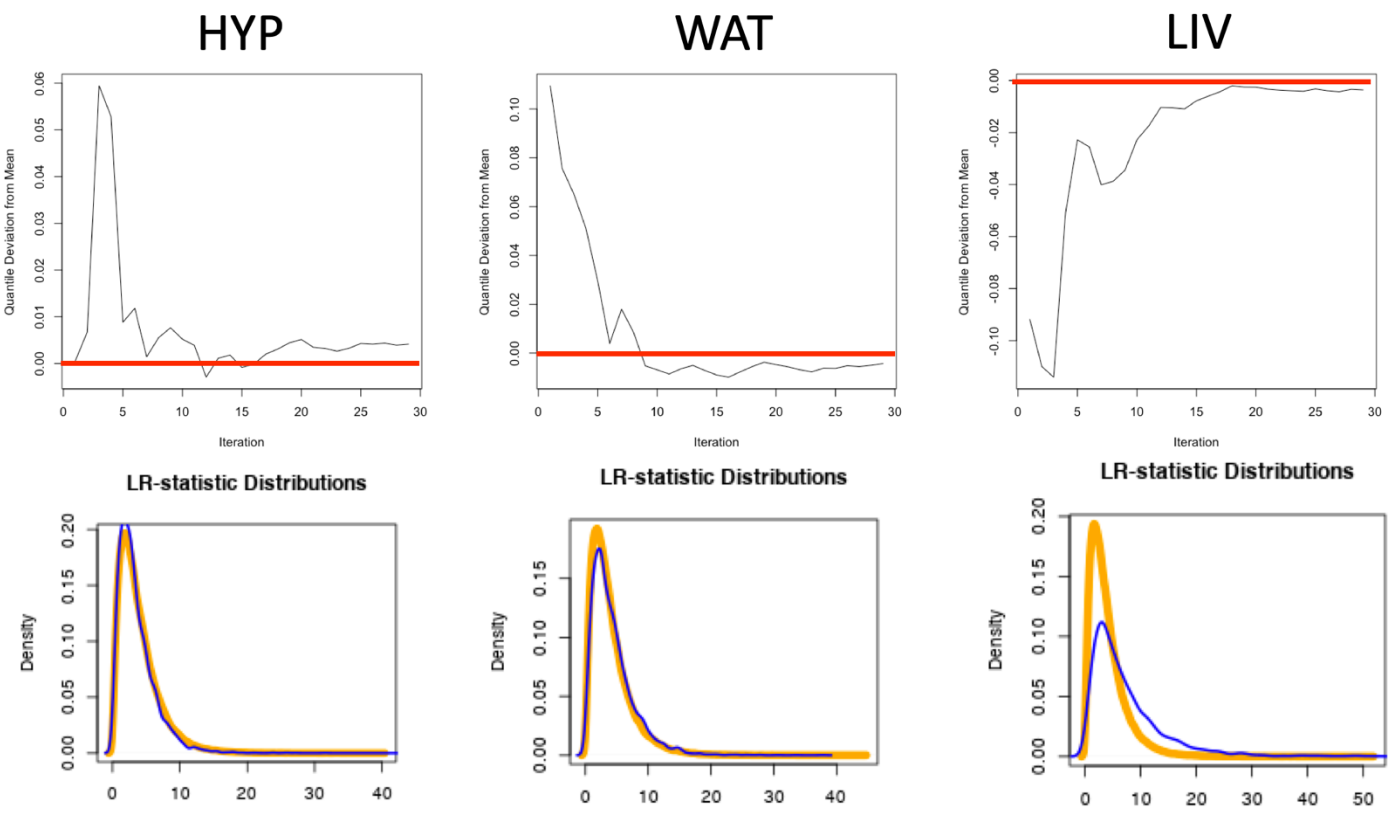


**Supplemental Figure 8: Stable null permutation plots for differential expression by cross.** Context was permuted to generate an appropriate null distribution of likelihood ratio statistics for each model. Null distribution stability was evaluated by calculating the total quantile deviation (TQD) at each iteration. The TQD of likelihood ratios under the null model approaches zero (red line) over thirty iterations. By thirty shuffles the TQD value fluctuates ~0.005 units per iteration in the most complex model, meaning that at worst LR estimates vary by ± 0.005, which corresponds to a p-value estimate error of ± $7.18x10^{-7}$ applying Wilk's Theorem to estimate significance from a $X^2$ distribution. The null distribution of likelihood ratio statistics (**Orange**) is plotted against the observed likelihood ratio distribution (**Blue**) for all tissues. Hyp = hypothalamus; WAT = white adipose; LIV = liver.

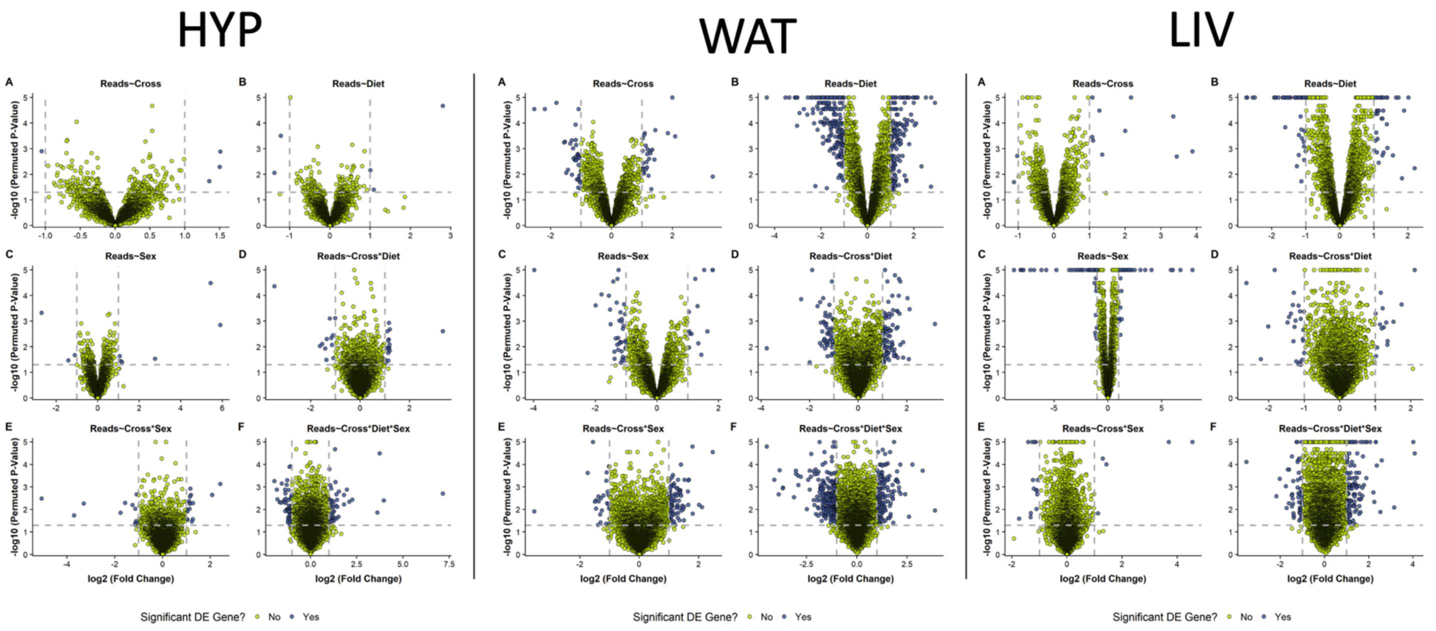


**Supplemental Figure 9: Volcano plots of differentially expressed genes.** $Log_{10}$ transformed permuted p-value plotted against $log_2$ transformed fold change for each context in each tissue. Hyp = hypothalamus; WAT = white adipose; LIV = liver.

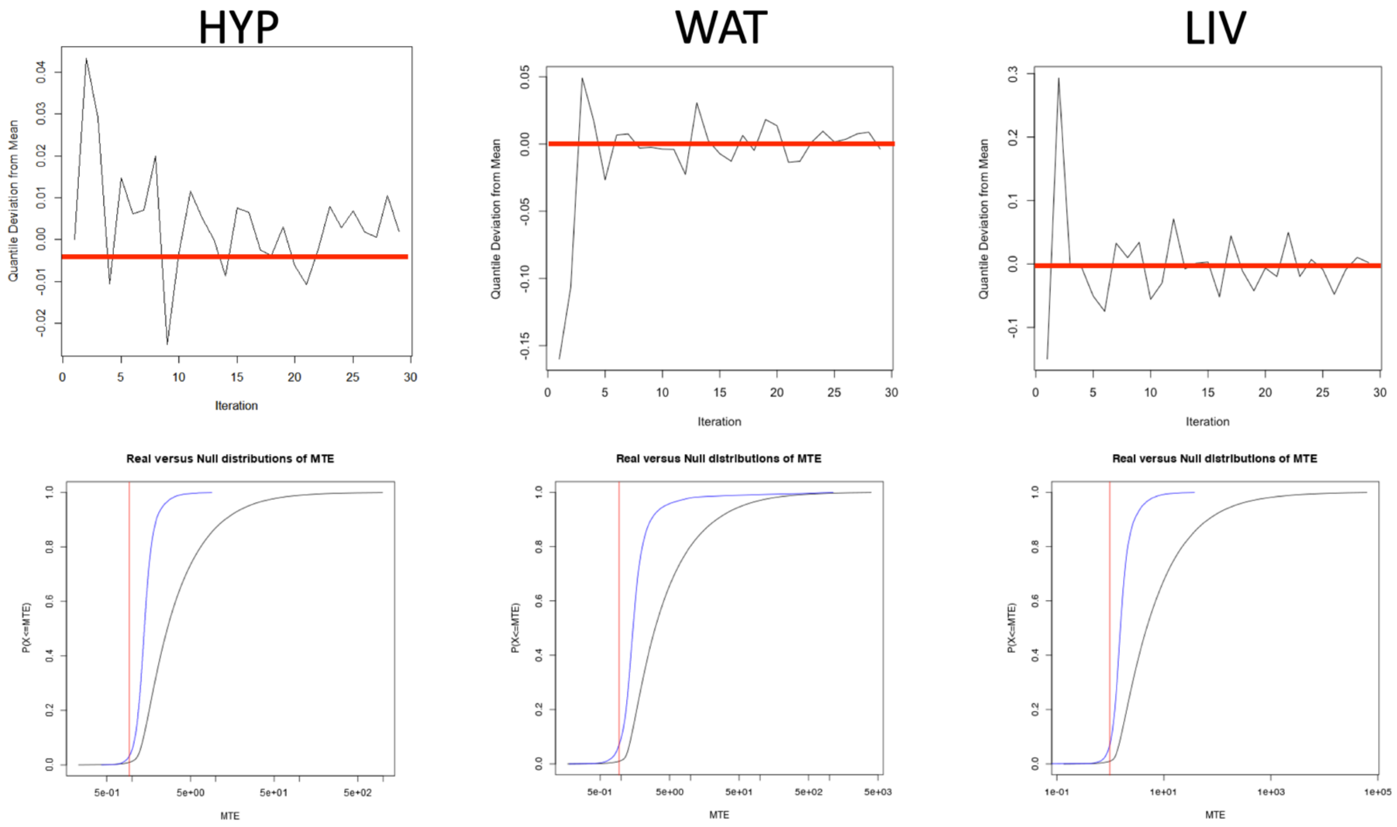


**Supplemental Figure 10: Stable null permutation plots for network pairs**. Context was permuted to generate an appropriate null distribution of mean test errors (MTE). Null distribution stability was evaluated by calculating the total quantile deviation (TQD) at each iteration. TQD is the sum of deviations of quantile values (Q) from the mean of quantile values of all prior iterations. Quantiles corresponded to one percent increments of the distribution. The TQD of MTE under the null model approaches zero (red line) over thirty iterations. By thirty shuffles the TQD value fluctuates ~0.01 units per iteration in the most complex model, meaning that at worst MTE estimates vary by ± 0.01, which corresponds to a predictive error of 1% of the standard deviation. The cumulative null distribution of MTE (**Black**) against which the observed likelihood ratio distributions can be compared (**Blue**). The alpha critical value for alpha=0.01 of the cumulative null distribution is indicated by the red vertical line. Hyp = hypothalamus; WAT = white adipose; LIV = liver.

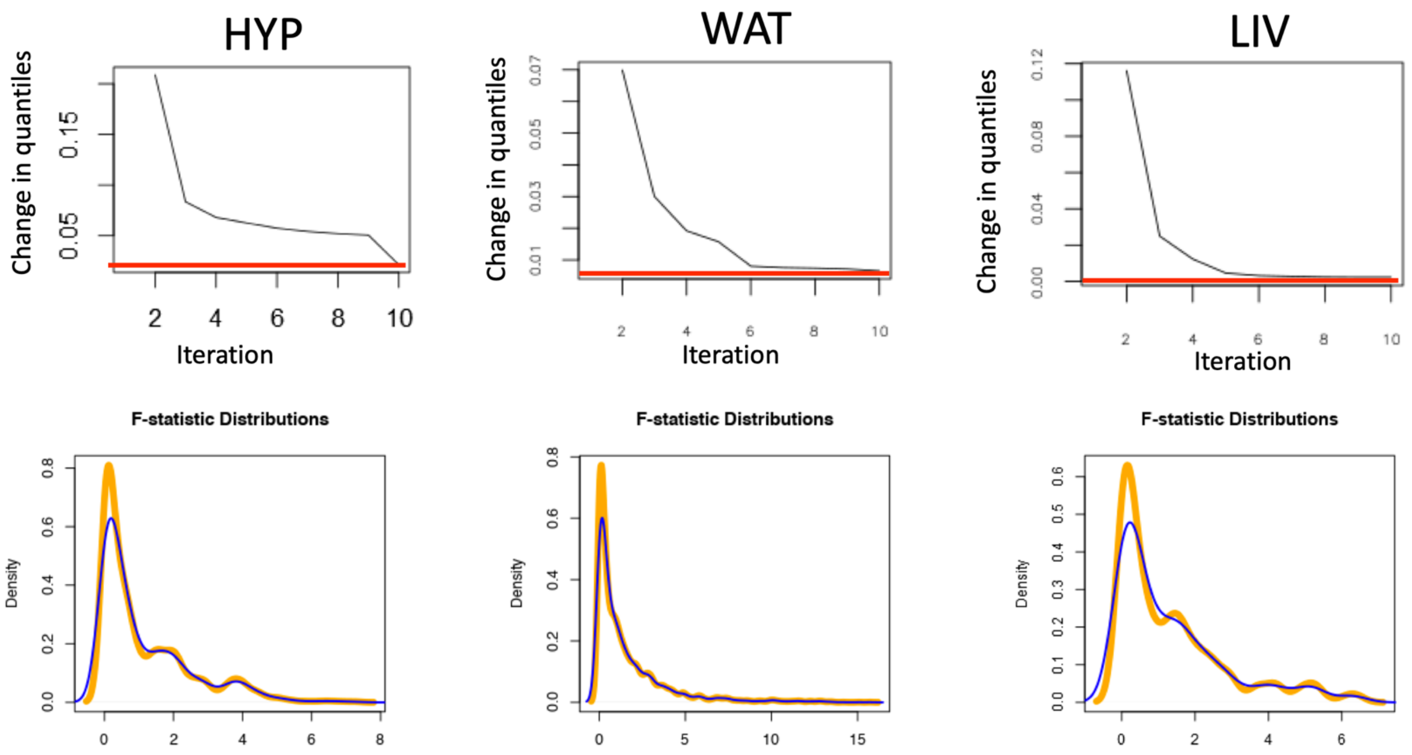


**Supplemental Figure 11: Stable null permutations plot for epistasis**. Imprinting scores were permuted to generate an appropriate null model of F-statistics. Null distribution stability was evaluated by calculating the total quantile deviation (TQD) at each iteration. The TQD of F-statistics under the null model approaches zero (red line) over ten iterations. By ten shuffles the TQD value fluctuates ~0.01 units per iteration, meaning that at worst F-statistics estimates vary by ± 0.01, which corresponds to a p-value estimate error of ± $6.71 \times 10^{-4}$ applying the standard F-distribution to estimate significance. The null distribution of F-statistics (**Orange**) is plotted against the observed F-statistic distribution (**Blue**) for all tissues. Hyp = hypothalamus; WAT = white adipose; LIV = liver.

**SupplementalTable1.xlsx : Allele-specific expression**
**SupplementalTable2.xlsx : Biallelic genes differentially expressed by cross**
**SupplementalTable3.xlsx : Networks of genes showing parent-of-origin allele-specific expression interacting with biallelic genes that are differentially expressed by cross**
**SupplementalTable4.xlsx : Over-representation input/output**
**SupplementalTable5.xlsx : Epistasis results**
**SupplementalTable6.xlsx : Alignment summaries**
**SupplementalTable7.xlsx : List of imprinted genes queried**
**SupplementalTable8.xlsx : QTL traits, positions, and genes falling within support intervals**
